# A global lipid map reveals host dependency factors conserved across SARS-CoV-2 variants

**DOI:** 10.1101/2022.02.14.480430

**Authors:** Scotland E. Farley, Jennifer E. Kyle, Hans C. Leier, Lisa M. Bramer, Jules Weinstein, Timothy A. Bates, Joon-Yong Lee, Thomas O. Metz, Carsten Schultz, Fikadu G. Tafesse

## Abstract

A comprehensive understanding of host dependency factors for SARS-CoV-2 remains elusive. We mapped alterations in host lipids following SARS-CoV-2 infection using nontargeted lipidomics. We found that SARS-CoV-2 rewires host lipid metabolism, altering 409 lipid species up to 64-fold relative to controls. We correlated these changes with viral protein activity by transfecting human cells with each viral protein and performing lipidomics. We found that lipid droplet plasticity is a key feature of infection and that viral propagation can be blocked by small-molecule glycerolipid biosynthesis inhibitors. We found that this inhibition was effective against the main variants of concern (alpha, beta, gamma, and delta), indicating that glycerolipid biosynthesis is a conserved host dependency factor that supports this evolving virus.

## Main Text

SARS-CoV-2 interacts with host membranes at every stage of its life cycle. It directly crosses the plasma membrane to enter the cell, replicates inside host-derived membrane compartments, acquires its envelope from the host, and traffics through the Golgi and lysosome to exit the cell. All viruses, by their nature, are wholly dependent on host pathways to meet their metabolic, structural, and trafficking needs, and to be effective, they must modulate these host pathways in some way. One dramatic example of this is the way in which SARS-CoV-2 re-engineers the host internal membranes into double-membraned vesicles (DMVs) and regions of convoluted membrane (CM) to facilitate its replication *(1, 2)*. This general pattern of membrane rearrangements is common among (+)-stranded RNA viruses *(3–5)*, although the specific structures vary by species. In flaviviruses such as Zika virus *(6)* and dengue virus *(7)*, these large-scale membrane alterations are accompanied by vast and varied changes at the molecular lipid level.

There are many preliminary lines of evidence suggesting that manipulation of host lipids may be a fundamental feature of SARS-CoV-2 infection. Several lipids and lipid-associated proteins have been identified as biomarkers of infection, including VLDL and HDL particles *(8)*, steroid hormones and various apolipoproteins *(9)*, while both elevated triacylglycerol (TAG) *(10)* and polyunsaturated free fatty acids *(11)* have been implicated as markers of severe disease outcomes. Furthermore, metabolic disorders such as obesity, diabetes, and hypertension have been described as key risk factors among patients who develop severe disease *(12)*. These observations indicate systemic changes in lipid metabolism at an organismal level, but it is still unknown how the virus alters the host lipid metabolism at a cellular level, and how these changes support the viral life cycle.

We hypothesized that SARS-CoV-2 would reprogram host lipid biosynthesis, and that the virus would depend on specific host metabolic pathways to survive and replicate effectively. To obtain a comprehensive understanding of how SARS-CoV-2 remodels the cellular lipid composition, we performed a detailed lipid survey of both infected cells and cells ectopically expressing individual SARS-CoV-2 proteins, assessing changes in host lipid composition as a result of infection and as a result of the activity of specific viral proteins. Based on our initial results, we examined lipid droplet flux during infection, and further interrogated the requirements for specific host lipids using small-molecule inhibitors of glycerolipid biosynthesis in multiple strains of SARS-CoV-2.

### Lipidomics of SARS-CoV-2 infected human cells

We performed global lipidomic profiling of HEK293T cells overexpressing the ACE2 protein, which we infected with SARS-CoV-2 or mock-infected for 24 hours (Fig 1A). Each condition was repeated in biological quintuplicate. Total cellular lipids were extracted following the method of Bligh-Dyer *(13)* and analyzed by liquid chromatograph electrospray ionization tandem mass spectrometry (LC-ESI-MS/MS). The abundances of the identified lipids were normalized by comparison to internal standards for quantitative analysis. In total, we identified 514 unique lipids spanning the glycerolipid, phospholipid, sphingolipid, and acylcarnitine categories (Supplementary Data 1). Of these, 409 (79.6%) were statistically altered between SARS-CoV-2 and mock infection (Benjamini-Hochberg adjusted p < 0.05, analysis of variance [ANOVA] test), changing between 2- and 64-fold in response to infection. Principal component analysis (PCA) of these observations confirmed that infection status accounted for most of the changes (Fig 1B), with the five infected samples and the five mock samples falling into two distinct groups.

**Fig. 1.**
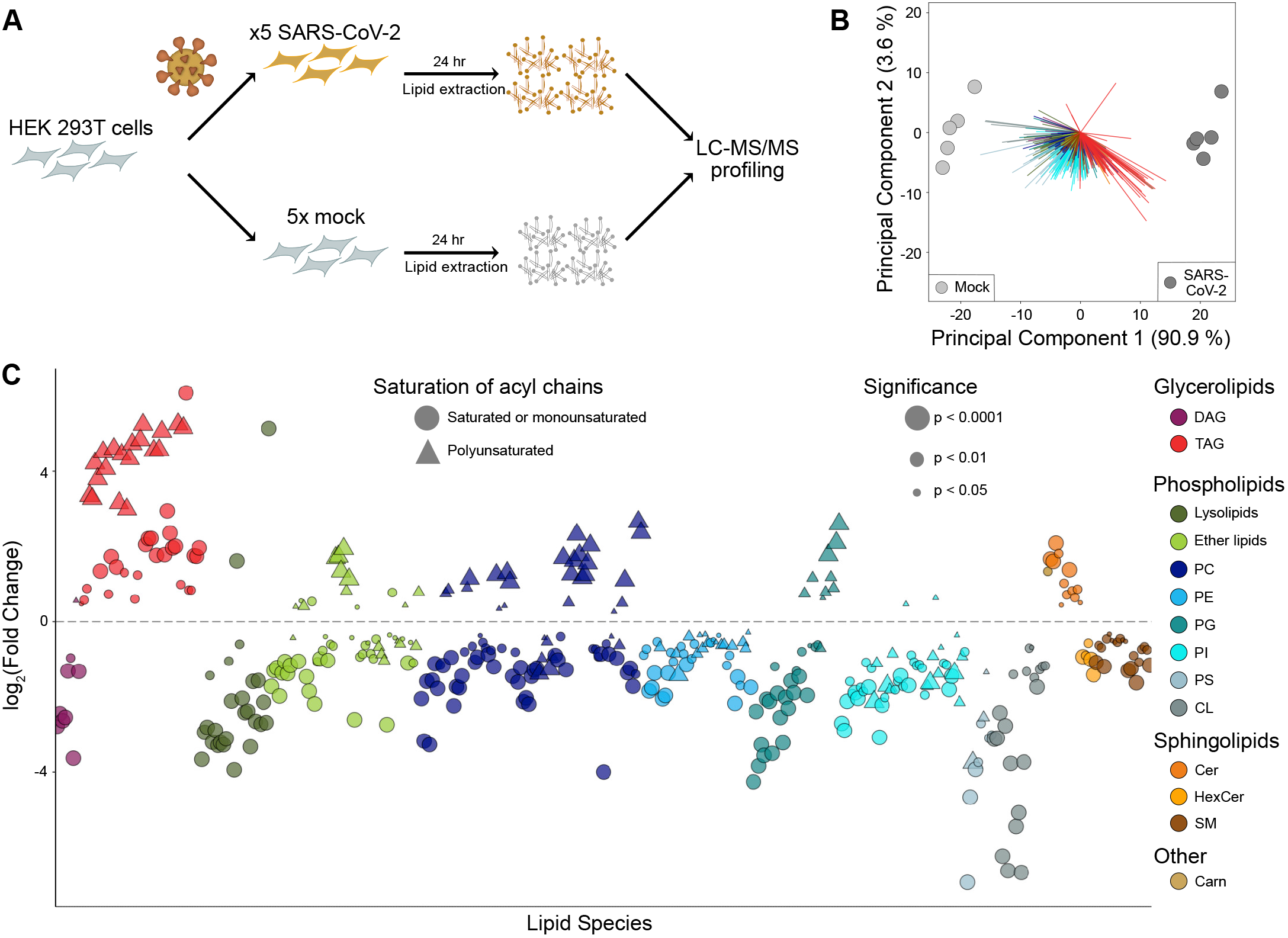
SARS-CoV-2 alters the lipid composition of its host cells. (**A**) Lipidomics study design. Each condition (SARS-CoV-2 infected or mock-infected HEK293T-ACE2 cells) is repeated in biological quintuplicate, and after 24 hours, cellular lipids are extracted and characterized by LC-MS/MS. (**B**) Principal component analysis of cells infected with SARS-CoV-2 or mock-infected. (n = 5 biological replicates, each point represents one biological replicate). (**C**) Individual lipid species characterized by abundance in SARS-CoV-2 infection relative to mock. Only significantly changed (p < 0.05, ANOVA, with Benjamini-Hochmini adjustment for multiple comparisons) lipids are shown. Log2(Fold Change) relative to mock infection is shown on the x-axis. Individual lipid species are colored by the class of lipid that they belong to. DAG = diacylglycerol; TAG = triacylglycerol; PC = phosphatidylcholine; PE = phosphatidylethanolamine; PG = phosphatidylglycerol; PI = phosphatidylinositol; PS = phosphatidylserine; CL = cardiolipin; Cer = ceramide; HexCer = hexosylceramide; SM = sphingomyelin; Carn = acylcarnitine.

We then examined how these changes in host lipid composition broke down based on class and acyl chain. Glycerolipids and phospholipids showed the largest and most significant changes (Fig 1C and Fig S1), with increasing triacylglycerol (TAG) and decreasing cardiolipin (CL) being the most altered. Examining the nature of the individual lipid species that changed in more detail (Fig 1C), we observed that the TAG species change based on their fatty acid composition. TAG species that bear polyunsaturated fatty acid (PUFA) chains were increased an average of 8-fold more than saturated or monounsaturated species. This trend was also observed in phospholipids: saturated phospholipids (phosphatidylcholine, PC; phosphatidylethanolamine, PE; phosphatidylglycerol, PG; phosphatidylinositol PI) almost universally decreased, while many polyunsaturated species increased, notably P-PC (phosphatidylcholine, plasmalogen-linked) by 2.7-fold, PC by 1.5-fold, and PG by 1.7-fold. Other notable phenotypes included a decrease in lysolipids (Lyso-PE by 5.1-fold and Lyso-PC by 3.1 fold), a decrease in CL by 5.9-fold, and an increase in ceramide (Cer) by 1.8-fold.

### Lipidomics of human cells ectopically expressing SARS-CoV-2 proteins

The genome of SARS-CoV-2 encodes 29 individual proteins (Fig 2A). Some of these proteins have been directly studied for SARS-CoV-2, but the roles of most of them must be extrapolated by comparison with the proteins of SARS-CoV, which are better studied. Several SARS-CoV proteins directly manipulate cellular membranes — nsp3, nsp4, and nsp6 together are known to induce DMVs *(14)* and CMs *(15, 16)* characteristic of coronavirus infection *(17, 18)*, and orf6 *(19)* also has a dramatic membrane-bending phenotype. Some proteins of SARS-CoV-2 — nsp1, nsp8, nsp9, nsp16 — have direct effects on mRNA splicing, or protein expression and membrane integration *(20)*. Many proteins of both viruses contain transmembrane domains (nsp2 *(21)*, orf7a *(22)*, orf7b *(23)*, orf3a *(24, 25)*) or lipid binding pockets (orf9b *(26)*) of unknown function, and many others, including nsp3 *(27)*, nsp6 *(28)*, orf3a *(29, 30)*, orf6 *(31)*, and orf7a *(32)*— mediate cell distress pathways such as apoptosis, autophagy, and the unfolded protein response (UPR), which are all known to alter cellular lipid composition *(33–35)* (Fig 2B).

**Fig. 2.**
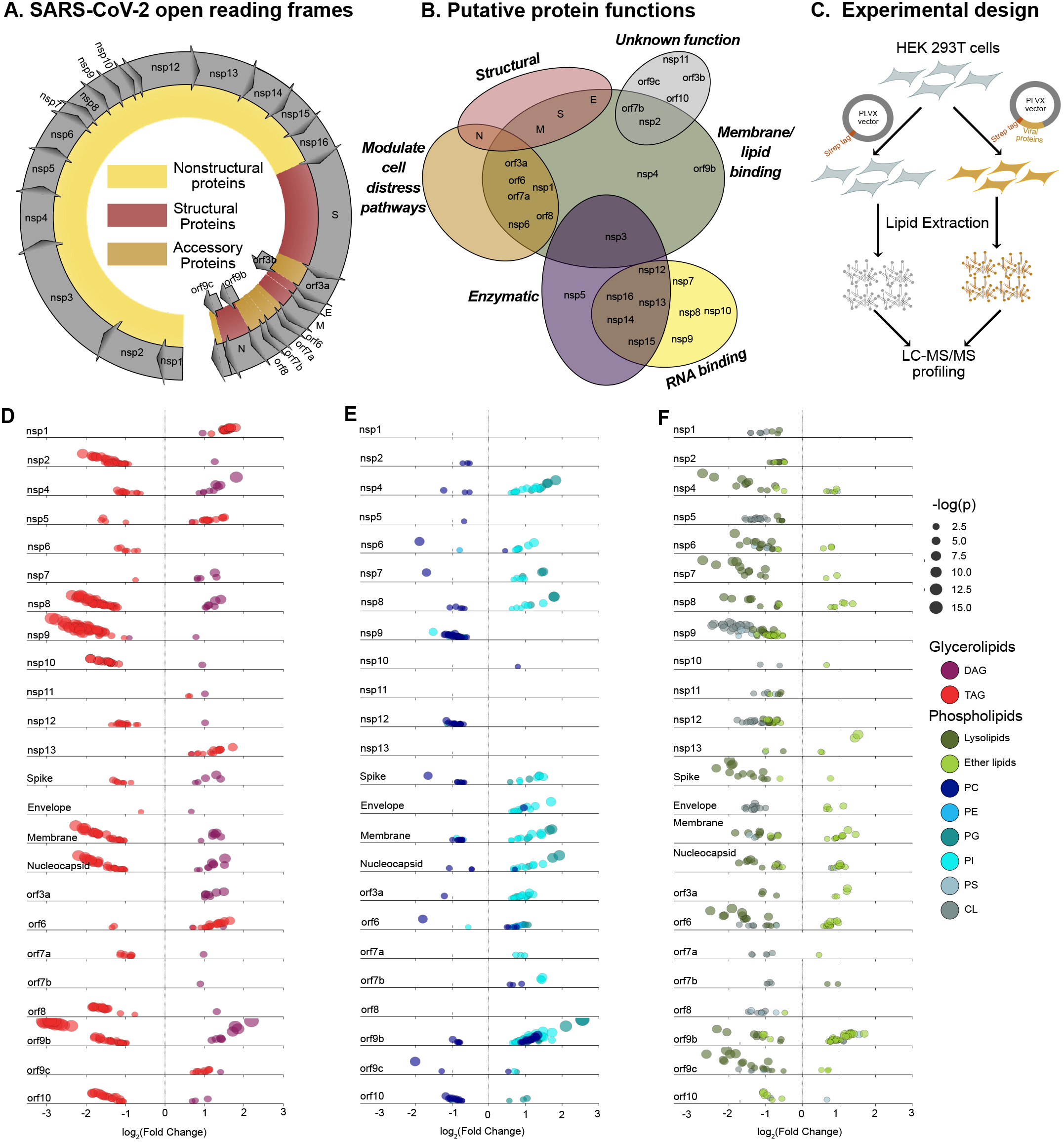
Ectopic expression of SARS-CoV-2 proteins modulates host lipid metabolism. **(A)** Open reading frames of the SARS-CoV-2 genome (**B**) Putative functions of SARS-CoV-2 proteins, based on early studies and sequence similarity to proteins from SARS-CoV. (**C**) Lipidomics study design. Each protein is transfected into HEK293T cells in biological quintuplicate, and after 48 hours, cellular lipids are extracted and characterized by LC-MS/MS. (**D-F**) Individual lipid species characterized by class and family. Only significantly changed (p < 0.05, ANOVA, Benjamini-Hochmini adjusted for multiple comparison) lipids are shown. Log2(Fold Change) relative to empty vector is shown on the x-axis. Individual lipid species are colored by the class of lipid that they belong to. Abbreviations same as Figure 1.

To assess how each SARS-CoV-2 protein affects host lipid metabolism, we performed untargeted lipidomic profiling of cells transfected with each viral protein, expressed in the PLVX vector with a C-terminal Strep tag. We optimized the expression of each protein in HEK-293T cells, measuring transfection efficiency by immunofluorescence of the Strep tag (Fig S2). In order to make meaningful comparisons between these conditions, high transfection efficiency was required (> 70%). Despite our efforts, this level of efficiency was not achieved for five proteins (nsp3, nsp14, nsp15, nsp16, orf3b); therefore, we continued on with the remaining 24. Each viral protein, as well as the PLVX empty vector, was used to transfect 6-cm dishes of HEK293T cells, in biological quintuplicate. After 48 hours of transfection, total cellular lipids were extracted following the method of Bligh-Dyer19 and analyzed by liquid chromatography electrospray ionization tandem mass spectrometry (LC-ESI-MS/MS) (Fig 2C). The abundances of the identified lipids were normalized by comparison to internal standards for quantitative analysis. In total, we identified 396 unique lipids spanning the glycerolipid, phospholipid, sphingolipid, and acyl-carnitine categories (Supplementary Data 2). Of these, 317 (80%) were significantly changed in at least one transfection (Benjamini-Hochberg adjusted p < 0.05, ANOVA).

For the samples transfected with viral proteins, we performed an EASE (Expression Analysis Systematic Explorer) score enrichment test of statistically significant lipids using Lipid MiniOn41. Lipid Mini-On performs enrichment analyses of lipidomics data using a text-mining process that bins individual lipid names into multiple lipid ontology groups based on their classification42 and other characteristics, such as chain length and number of double bonds. Using Lipid Mini-On we found that the most common enrichments that were increased with the viral protein transfections were PIs, (elevated in 21 of 28 transfections), diacylglycerols (DAGs) (12 transfections), and ether-linked lipids, in particular vinyl-ether phosphatidylcholines (O-PC) (10-12 transfections), Cer (10 transfections), and TAG (6 transfections). Enrichments that were found to be decreased were Lyso-PC (decreased in 21 transfections), CLs (12 transfections), which almost universally decrease in abundance, and TAGs (decreased in 14 transfections). (Summarized in Fig 2, D-F; fold changes for all significant lipids are shown in Fig S3.) The 24 viral proteins studied show a wide variety of lipid alterations, suggesting that SARS-CoV-2 influences host lipid metabolism in diverse ways through multiple molecular mechanisms.

### Correlating live virus and viral protein lipidomic phenotypes

With two substantial datasets of virus-induced lipid changes, we sought to link the changes observed in live virus infection to the action of specific viral proteins. First, we performed unsupervised clustering of the normalized lipid species observed in the protein-transfected dataset by t-SNE (Fig 3A). While most phospholipids did not cluster substantially, TAG, in particular, formed distinct clusters, and, in an echo of the live virus phenotype, saturated species and poly-unsaturated species clustered separately. Of note, two other molecular features of infection — Cer and CL — also sorted into distinct clusters.

**Fig. 3.**
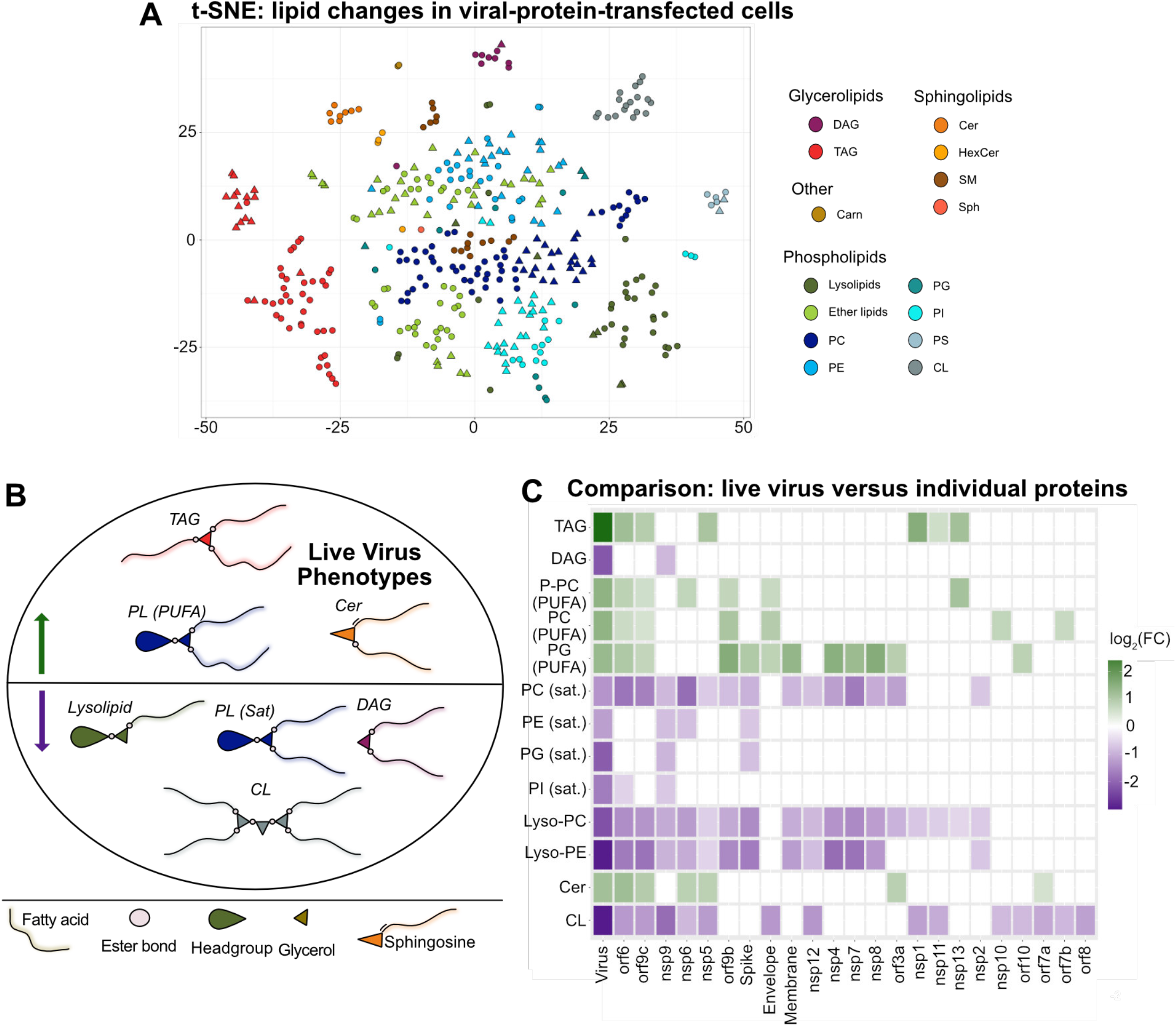
Individual SARS-CoV-2 proteins recapitulate overlapping lipid features of live infection. **(A**) Unsupervised clustering of the normalized lipid species observed in the protein-transfected dataset by t-SNE. Abbreviations same as Figure 1. (**B**) Summary of lipids altered upon infection with SARS-CoV-2. Cer = ceramide; PL (PUFA) = phospholipids bearing polyunsaturated acyl chains; TAG = triacylglycerol; PL (Sat) = phospholipids bearing saturated or monounsaturated acyl chains; CL = cardiolipin (**C**) Average fold change within each class described above in each condition, both live virus infection and ectopic protein expression. Only significantly changed (p < 0.05, see Fig 1 and Fig 2 for descriptions of statistical tests in the live virus and transfection conditions, respectively) lipid species were used in this calculation.

The dominant features of lipid remodeling in live virus infection were an increase in TAG, Cer, and phospholipids bearing polyunsaturated fatty acyl chains; and a decrease in lysolipids, DAG, CL, and saturated phospholipids (Fig 3B). In order to assess each viral protein for its ability to produce these changes, the average fold change for each of these classes was calculated for each condition (Fig 3C). Once again we saw that the virus has multiple proteins that influence remodeling of the lipid environment of its host cells, suggesting a distinct role for each viral protein. Each feature of infection was recapitulated by at least one protein, and different proteins appear to be responsible for different aspects of the live virus lipid phenotype.

In particular, TAG increase was recapitulated by six proteins (orf6, nsp13, nsp5, orf9c, nsp1, and nsp11). Cer increase was recapitulated by six as well (nsp6, orf6, nsp5, orf9c, orf3a, and orf7a), and polyunsaturated PC (both ether- and ester-linked) increase was recapitulated by four (orf6, orf9c, orf9b, and E). Of note, orf6 and orf9c recapitulated all three of these distinctive alterations, and also recapitulated the most individual phenotypes of any protein.

### Lipid droplet dynamics in SARS-CoV-2 infection

TAG is the most significantly and the most substantially increased lipid in response to viral infection. TAG is produced through the acylation of DAG by DGAT1 or DGAT2, where it is then sequestered in lipid droplets that can be accessed as a source of fatty acids. TAG breakdown is the result of several lipases that remove an acyl chain to produce DAG (Fig 4A). Lipid droplets (LDs) are the cellular reservoir for TAGs, and have well-established roles in the life cycles of other viruses. Hepatitis C virus (HCV) and rotaviruses both cause LDs to accumulate during infection, and HCV uses LDs as the site of viral assembly while rotavirus replication occurs in close proximity to lipid droplets *(36, 37)*. Dengue virus, meanwhile, consumes host lipid droplets and appears to use them as a source for beta-oxidation *(38)*.

**Fig. 4.**
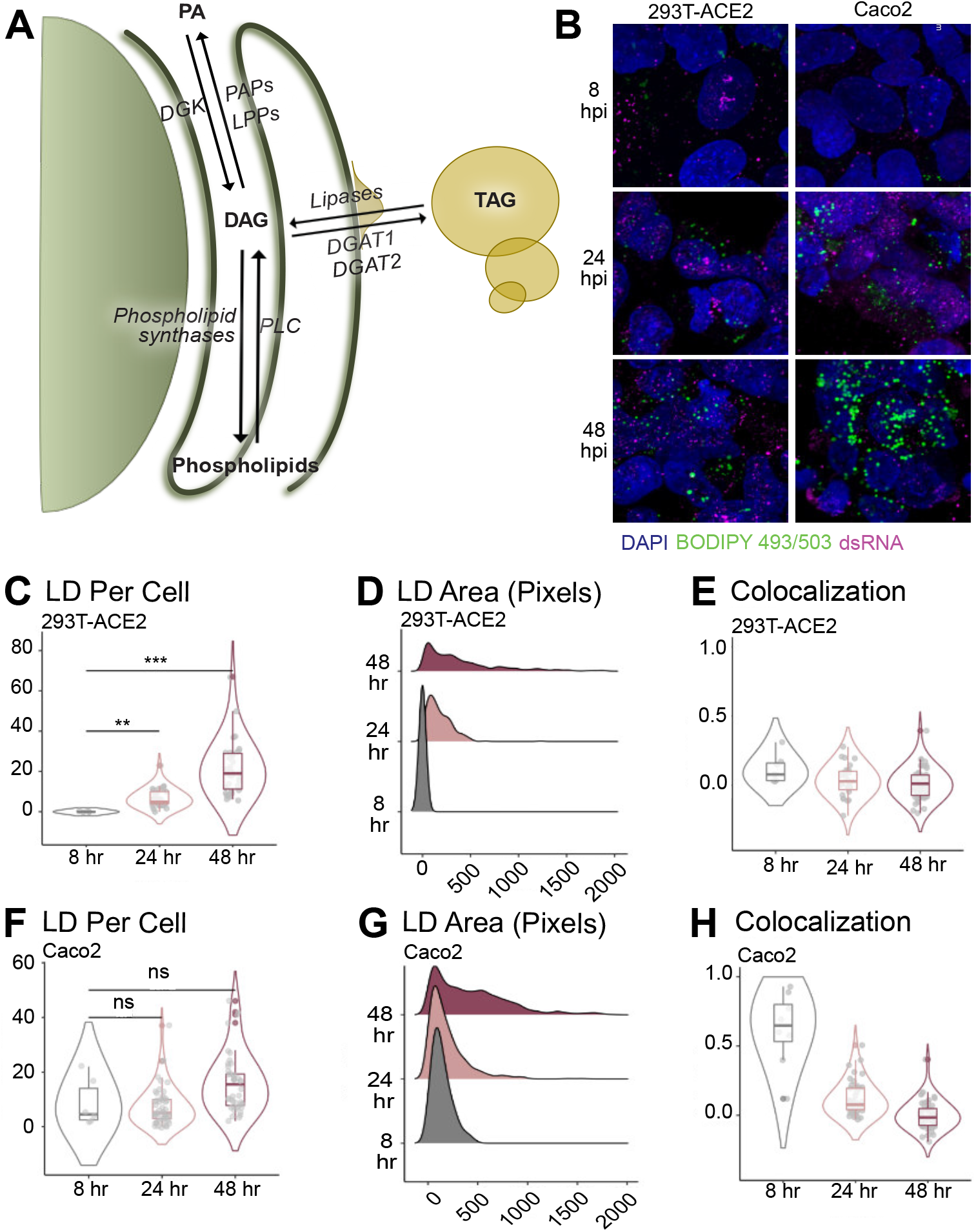
Lipid droplets are induced following SARS-CoV-2 infection and after the transfection of key viral proteins. (**A**) Overview of central glycerolipid metabolism. PA = phosphatidic acid; PAP = phosphatidic acid phosphatase; LPP = lysophosphatidic acid phosphatase; DGK: diacylglycerol kinase; DAG = diacylglycerol; TAG = triacylglycerol; DGAT 1/2 = diacylglycerol acetyltransferase 1/2; PLC = phospholipase C. (**B**) 293T-ACE2 and Caco-2 cells infected with SARS-CoV-2 strain USA-WA1/2020 (MOI = 1) and fixed at the indicated timepoints. LDs and infected cells were visualized with BODIPY 493/503 and anti-dsRNA immunofluorescence, respectively. Images are representative of three independent experiments. (**C & F**) Number of lipid droplets per cell; each data point is a cell. *p ≤ 0.05, **p ≤ 0.01, ***p ≤ 0.001, ***p < 0.0001, one-way ANOVA. (**D & G**) Distribution of the area of each lipid droplet, in pixels. (**E & H**) Colocalization of dsRNA and BODIPY by Pearson’s coefficient. Each data point is a cell. *p ≤ 0.05, **p ≤ 0.01, ***p ≤ 0.001, ***p < 0.0001, one-way ANOVA.

We sought to understand how the abundance and morphology of host lipid droplets changes during the course of SARS-CoV-2 infection, and whether they are associated with virus-induced membrane structures. We chose BODIPY 493/503, a bright, hydrophobic dye, to mark the lipid droplets, a well-established method*(39)*. We also used an anti-dsRNA antibody to mark the sites of viral replication; dsRNA is an intermediate in the synthesis of the virus’s RNA genome, and has been shown to localize to DMVs*(14)*. We visualized both of these markers 8 hours, 24 hours, or 48 hours post infection in HEK-293T cells and then stained with BODIPY 493/503 to mark lipid droplets and an anti-dsRNA antibody to mark the site of viral replication (Fig 4B). We see a clear increase in the number and size of lipid droplets in a time-dependent manner over the course of infection, quantified in Fig 4C-D. Lipid droplets per cell increase from zero at 8hpi, to an average of 6.7 at 24hpi, to an average of 21.5 at 48hpi. Lipid droplet area increases from zero pixels per droplet at 8hpi, to an average of 177 pixels per droplet at 24hpi, to an average of 400 pixels per droplet at 48hpi. However, there does not appear to be any colocalization of the lipid droplets and dsRNA, suggesting that the virus is not using lipid droplets directly as a platform for replication (Fig 4E).

To further validate these observations, similar experiments were performed in the human epithelial Caco2 cell line. Here, a slight increase in lipid droplet number was observed, from an average of 8 lipid dropletss per cell at 8 and 24hpi, to an average of 15.9 lipid droplets per cell at 48hpi, although the increase was not significant. Lipid droplet area, however, did significantly increase throughout the course of infection, to a similar degree as in HEK293T-ACE2 cells, from an average of 136.5 pixels per droplet at 8hpi to an average of 192.5 pixels per droplet at 24hpi to an average of 431.1 pixels per droplet at 48hpi (Fig 4F-G). Once again, colocalization with dsRNA was not observed (Fig 4H).

### Viral requirements for central glycerolipid metabolism

Since levels of individual glycerolipid species as well as glycerolipid-based structures were altered by infection, we asked whether these pathways are necessary for viral proliferation. We selected an array of commercially available small molecule inhibitors of lipid synthesis, focusing on inhibitors of de novo neutral lipid synthesis as well as lipolytic enzymes of lipid recycling (Fig 5A-F). We performed initial cytotoxicity measurements using a resazurin-based viability assay *(40)* (Fig S4) and selected a non-cytotoxic concentration of each compound to screen for inhibition of viral infection. 293T-ACE2 cells were treated overnight with each compound, and then infected with SARS-CoV-2. After 48 hours of infection, culture supernatants were collected and the amount of infectious virus produced in the presence of each compound was quantified by focus forming assay *(41)*.

**Fig 5.**
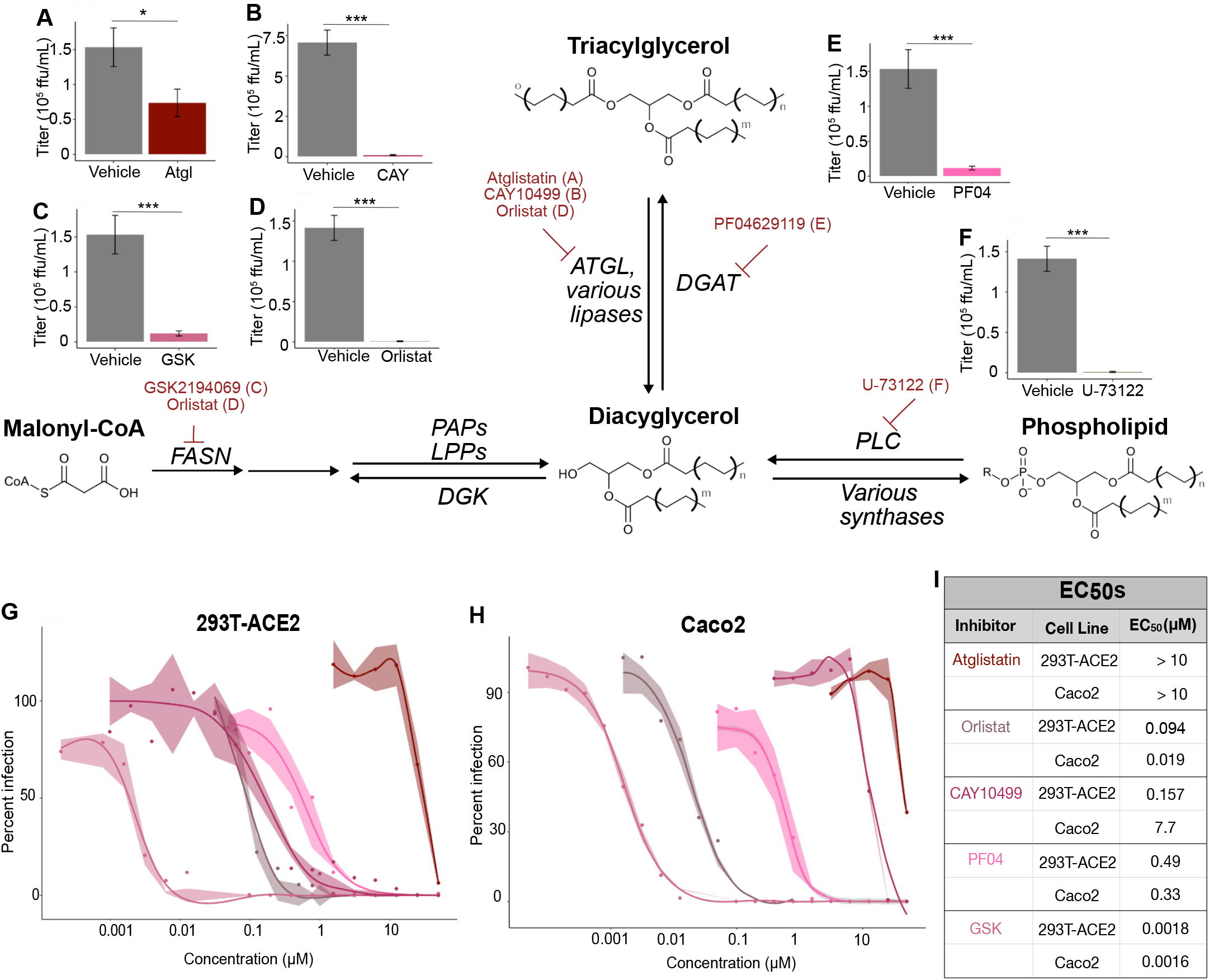
Central glycerolipid metabolism is essential for SARS-CoV-2 infection. (**A-F**) Screen of neutral lipid biosynthesis inhibitors. HEK-293T-ACE2 cells were treated with 10 μM of each compound overnight prior to infection. Cells were infected for 48 hours prior to supernatant collection and focus forming assay. Bars represent viral titers from cells treated with the indicated inhibitors, measured by focus forming assay. Data are mean ± SE; *p ≤ 0.05, **p ≤ 0.01, ***p ≤ 0.001, ***p < 0.0001, one-way ANOVA. Data are from three independent experiments. FASN = fatty acid synthase; PAP = phosphatidic acid phosphatase; LPP = lipid phosphate phosphatase; DGK = diacylglycerol kinase; ATGL = adipose triacylglycerol lipase; DGAT = diacylglycerol acetyltransferase; PLC = phospholipase C (**G**) EC_50_ curves for selected inhibitors in 293T-ACE2 cells. HEK-293T-ACE2 cells were treated with 2-fold dilutions of each compound overnight prior to infection. Cells were infected for 48 hours prior to supernatant collection and focus forming assay. Percent infection is calculated as [Titer(inhibitor) /Titer(vehicle)]*100. Data are from three independent experiments. Shadow is SE. Curve fits are calculated using a nonlinear curve fit to the Hill equation: Response = (Max Response)/(1 + [EC_50_/Concentration]^n), where the max response is defined as 100% inhibition. (**H**) EC_50_ curves for selected inhibitors in Caco2 cells. Experiment and analysis same as described in (G) (**I**) EC_50_ values from the curves in G and H. EC50 values are calculated from the curve fit described above.

This screen revealed several steps of lipid biosynthesis which are essential to the production of infectious virions. De novo fatty acid synthesis appeared critical, as GSK2194069, an inhibitor of fatty acid synthase (FASN) *(42)*, as well as orlistat, a nonspecific lipase inhibitor and inhibitor of fatty acid synthetase FASN *(43, 44)*, an FDA-approved drug, both completely blocked viral production (Fig5C, D). TAG synthesis and lipolysis are both required, as PF-04620110, an inhibitor of DGAT1 *(45)*, Orlistat, and CAY10499, which is a non-specific lipase inhibitor *(46, 47)*, all blocked infection (Fig 5E, D, B). Atglistatin *(48)*, which specifically blocks adipose triacylglycerol lipase, partially inhibited viral production (Fig 5A), suggesting that broad-spectrum lipase inhibition is more effective than inhibiting only one lipase. The importance of DAG production to the virus, perhaps as a precursor to TAG, is indicated by the efficacy of U-73122 (Fig 5F), which inhibits phospholipase-C-dependent processes *(49)*.

To directly compare the inhibitors of central glycerolipid metabolism, we designed a more detailed study to test a range of concentrations for each inhibitor. We tested a range of two-fold dilutions of each compound, and in parallel with the focus-forming assay to assess viral replication, we performed a resazurin-based cytotoxicity assay to verify that any deficiency in viral production was not due to impaired cell viability (Fig S4). The most effective inhibitor by about fifty-fold was GSK2194069 (EC50 = 1.8 nM, 293T-ACE2). GSK2194069 blocks FASN, suggesting that de novo lipid synthesis is strictly required for viral survival. Orlistat followed in efficacy (EC50 = 94 nM, 293T-ACE2), highlighting the importance of both fatty acid synthesis and lipolysis to the virus. The other broad-spectrum lipase inhibitor, CAY10499 (EC50 = 157 nM, 283T-ACE2) had a similar efficacy to PF04620110 (EC50 = 490 nM, 293T-ACE2). Atglistatin, the most specific lipase inhibitor, became cytotoxic before complete inhibition was achieved, and so an EC50 could not be calculated; certainly it is higher than 10 μM, showing again that the virus is not dependent on the activity of one specific lipase, but rather on a certain lipid composition. Taken together, these results indicate a profound dependence on host lipid metabolism, and in particular glycerolipid flux. The de novo synthesis of TAG is required, as is the ability to release the fatty acids sequestered in this neutral storage lipid through lipolysis.

### Glycerolipid biosynthesis is a conserved host dependency factor for variants of SARS-CoV-2

Given that our most effective inhibitors all relate in some way to the dynamics of TAG production, we hypothesized that their efficacy is due to the virus’s specific requirements for lipid droplets. We performed microscopy of cells treated with selected inhibitors at 10 μM overnight (Fig 6A, quantified in Fig 6B, experimental scheme in Fig S5A). We once again observed that virus alone induced a significant increase in the number of lipid droplets per cell, from an average of 0.3 to average of 3, and further noted that in the absence of virus, none of the inhibitors had an effect on lipid droplet numbers. In the presence of virus, GSK2194069 treatment did not prevent a statistically-significant increase in lipid droplet numbers, while PF04620110 did, suggesting that DGAT1 is essential for virus-induced lipid droplet production. Orlistat, meanwhile, resulted in an increase in lipid droplet numbers relative to vehicle treatment during infection, from an average of 3 to an average of 7.5. These results underscore the specificity of SARS-CoV-2’s requirements for lipid droplets: while SARS-CoV-2 infection results in an overall increase in the lipid droplets in each infected cell, both TAG synthesis and lipolysis are required to support the production of infectious virions. Furthermore, simply increasing the number of lipid droplets does not support replication: pure accumulation of TAG resulting from the inhibition of lipolysis is as detrimental to infection as preventing its synthesis.

**Fig 6.**
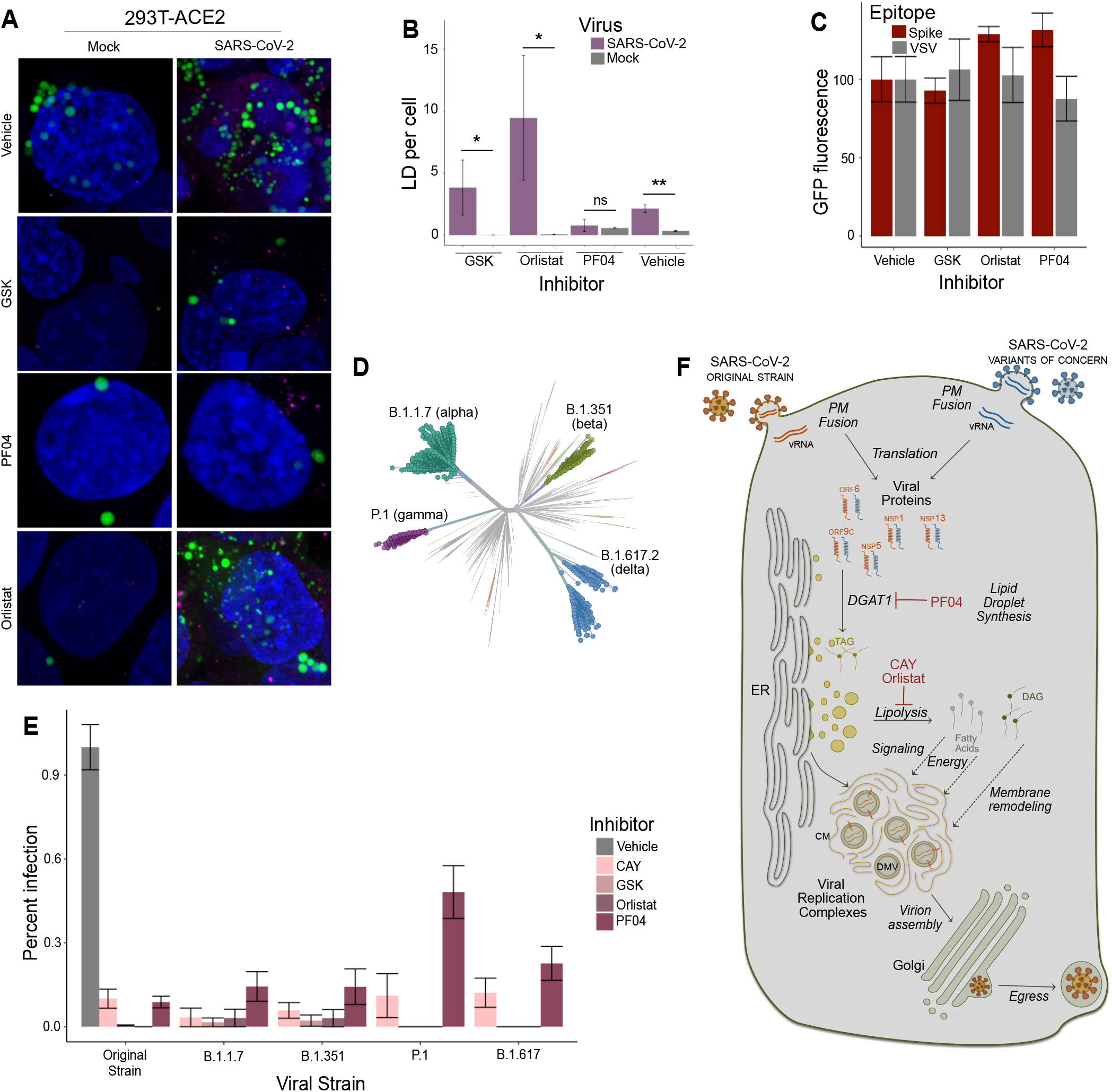
Mechanisms and breadth of glycerolipid inhibition against SARS-CoV-2. **(A)** Representative images of HEK293T-ACE2 cells treated with each indicated inhibitor (10 μM) or vehicle (DMSO), infected with SARS-CoV-2 (MOI = 1), and stained to visualize lipid droplets (BODIPY 493/503), and dsRNA. Images are representative of three independent experiments. (**B**) Quantification of lipid droplet numbers in (A). Data are mean ± SE; * p < 0.05, ** p < 0.01, *** < 0.001, one-way ANOVA (**C**) GFP fluorescence resulting from an infection with lentivirus pseudotyped with either SARS-CoV-2 Spike protein or VSV G protein. Estimated area of DAPI and GFP fluorescent pixels was calculated with built-in BZ-X software (Keyence) and GFP fluorescence was normalized to the DAPI signal for each condition. There were five biological replicates for each condition, and the biggest outlier was removed from analysis due to inherent variability in the assay. Data are mean ± SD. (**D**) A model for neutral lipid flux during SARS-CoV-2 infection. All SARS-CoV-2 genomes enter the cytosol (shown here by direct fusion with the plasma membrane; endosomal entry has also been reported, especially in Vero E6 cells, but is thought to be less physiologically relevant to human infection). Viral proteins are expressed by host metabolic machinery; orf6, orf9c, nsp1, nsp5, and nsp13 all directly induce TAG formation via DGAT1, which is inhibited by PF04620110. Lipid droplets proliferate following infection, and are also sources for raw lipid material released by lipolysis, which is inhibited by CAY10499 and orlistat. These raw materials may be sources of energy, signaling mediators, and lipids for the creation of viral replication complexes. Assembled virions are trafficked through the Golgi and released from the cell by lysosomal exocytosis. (**E**) Unrooted phylogenetic tree of SARS-CoV-2 variants of concern, generated by Nextstrain *(68,69)*, an open-source repository of pathogen genomic data. (**F**) Inhibition of the original strain and four variants of concern of SARS-CoV-2 in 293T-ACE2 cells by four inhibitors of glycerolipid biosynthesis, each at 10 μM overnight prior to an 48-hour infection. Data are from three independent experiments; data are mean ± SE.

SARS-CoV-2 interacts with host lipids at every stage of its life cycle. To rule out the possibility that glycerolipid metabolism is necessary for the initial attachment and endocytosis of the virus, we performed an entry assay using S-pseudotyped lentivirus. For this experiment, lentiviruses were generated that display the SARS-CoV-2 S protein and carry a GFP reporter; lentiviruses coated instead with the VSV G protein were used as a control. Successfully infected cells express GFP, and quantitative microscopy was used to assess infection (Fig S5B). 293T-ACE2 cells were treated overnight with selected inhibitors of glycerolipid biosynthesis and then infected with either of these two lentivirus constructs. We did not observe a significant reduction in viral entry in the presence of any of the inhibitors tested, suggesting that the virus depends upon this lipid biosynthetic pathway to facilitate the intracellular stages of its life cycle (Fig 6C).

The continued global transmission of SARS-CoV-2 has led to the emergence of variants of concern (VOC) that show evidence of increased transmissibility *(50)*or resistance to prior immunity *(51, 52)* (Fig 6D). The major VOCs include the B.1.1.7 (also called the alpha variant), first identified in southeast England in November 2020 *(53)*; B.1.351 (beta variant), identified in November 2020 in South Africa *(54)*; P.1 (gamma variant), identified in December 2020 in Brazil *(55)*; and B.1.617.2 (delta variant), identified in October 2020 in India *(56)*. Several recent studies have shown that these strains escape neutralization of serum antibodies collected from individuals that received COVID-19 vaccine or were previously infected. Most of the mutations in the emerging VOCs are on the spike protein, and while there are some reported alterations in non-structural proteins, mutations that fundamentally perturb the virus’s ability to manipulate host pathways likely come with a quite high fitness cost. We hypothesized, therefore, that the replication of the variants of SARS-CoV-2 is inhibited to a similar degree to the original USA/WA1/2020 strain.

To test if the small molecules that inhibit glycerolipid biosynthetic machinery are broadly efficacious, we used the P.1, B.1.351, B.1.1.7, and B1.617.2 strains, as well as the WA1 original strain, to infect cells that had been pre-treated overnight with 10 μM CAY10499, GSK2194069, PF04620110, and Orlistat, and assessed viral proliferation by focus forming assay. We performed these experiments in both 293T-ACE2 cells and Caco2 cells. We observed very few differences in efficacy of the compounds among the four strains tested (Figure 6E and Fig S6). GSK2194069 and Orlistat comprehensively block infection (< 5% of vehicle treatment) in both cell types and all five strains. CAY10499 has slightly different efficacies between the two cell lines, (~ 5-10% infection in HEK293T-ACE2, ~ 30% infection in Caco2), but there is no statistical difference between the variants within each cell line. PF04620110 resembles CAY10499 in Caco2 cells; in HEK293T cells, PF04620110 shows reduced efficacy against the P.1 strain. In the delta strain, CAY10499 showed a slightly significant reduction in foci in Caco2 cells (p = 0.045), from ~ 30% infection in WA to ~ 5% infection in delta; no other inhibitors were significantly different. Overall, these results show an encouraging conservation of inhibitor efficacy against the four variants of concern in two cell lines.

## Discussion

Based on our integrated lipidomics, microscopy and small-molecule inhibition experiments, we propose here a model for how SARS-CoV-2 uses lipid droplets to support infection (Figure 6F). We show that lipid droplet proliferation is a consequence of infection, and that both TAG synthesis and lipolysis are required for effective replication. The lipid droplet phenotype appears to be part of a profound reprogramming of cellular lipid metabolism which is induced directly by individual viral proteins; strikingly, polyunsaturated lipids are dramatically increased while saturated lipids are decreased, suggesting that viral membrane structures require a particularly high level of fluidity. While lipid droplets do not appear to be parts of the viral replication complex, given the very low levels of colocalization between dsRNA and BODIPY in infected cells, it seems likely that their roles in buffering lipid levels and facilitating membrane plasticity support the ambitious coronaviral membrane rearrangements.

Using small-molecule inhibitors of glycerolipid metabolism, we showed that SARS-CoV-2 fundamentally requires host lipid metabolic pathways for its survival and proliferation. Our findings highlight the dynamic and specific involvement of host lipids in infection: SARS-CoV-2 requires both de novo fatty acid and TAG synthesis, and lipolysis, simultaneously promoting lipid synthesis and providing specific lipids for viral processes. We further showed that these inhibitors work as effectively against the recently emerging SARS-CoV-2 variants of concern as they do against the original WA1 strain, demonstrating the advantage of designing host-targeted therapeutics against a conserved host dependency pathway.

Our findings fill an important gap in our understanding of host dependency factors of coronavirus infection. Our systematic analysis of the protein-by-protein effect on host lipids reveals a complex network of many individual viral proteins responsible for diverse aspects of host lipid remodeling. Both of our lipidomics datasets are resources for understanding cellular disease pathology and suggest potential directions for therapeutic discovery, highlighted by the success of several inhibitors of glycerolipid biosynthesis in blocking viral replication. In light of the evolving nature of SARS-CoV-2, it is critical that we understand the basic biology of its life cycle in order to illuminate additional avenues for protection and therapy against this global pandemic pathogen, which spreads quickly and mutates with ease.

## Materials and Methods

### Materials

#### Cell lines

Cell lines (HEK293T, HEK293T-ACE2, Vero-E6, and Caco2) were obtained from ATCC.

#### Viral strains

SARS-CoV-2 viral strains (isolate USA-WA1/2020: Identifier #NR-52281; isolate USA/CA_CDC_5574/2020: Identifier #NR-54011; isolate hCoV-19/South Africa/KRISP-K005325/2020: Identifier #NR-54009; hCoV-19/Japan/TY7-503/2021: Identifier #NR54982; isolate hCoV-19/USA/PHC658/2021: Identifier # NR-55611) were obtained from BEI resources and propagated in Vero E6 cells.

#### Recombinant DNA

Plasmids containing strep-tagged SARS-CoV-2 proteins were obtained from the Krogon lab at UCSF (*59*).

#### Chemicals and antibodies

Inhibitors of lipid biosynthesis were obtained from Cayman Chemical; EquiSPLASH lipidomics internal standard was obtained from Avanti Polar Lipids. Anti-dsRNA antibody was obtained from Millipore (identifier MABE1134); anti-mouse IgG AlexaFluor 647 was obtained from Invitrogen (Identifier A32628); anti-llama secondary HRP, goat IgG was obtained from Novus (identifier NB7242).

### Methods

#### Cell culture

Unless otherwise stated, cells were maintained at all times in standard tissue culture-treated vessels in DMEM supplemented with 1% nonessential amino acids and 1% penicillin-streptomycin at 37 °C and 5% CO2. Media for Vero-E6 cells, 293T (wt) and 293T-ACE2 cells was supplemented with 10% FBS while media for Caco2 cells was supplemented with 20% FBS.

#### SARS-CoV-2 growth and titration

All SARS-CoV-2 isolates were obtained from BEI resources: USA/WA1/2020 (NR-52281), USA/CA CDC 5574/2020 [lineage B.1.1.7] (NR-54011), hCoV-19/South Africa/KRISP-K005325/2020 [lineage B.1.351] (NR-54009), hCoV-19/Japan/TY7-503/2021 [linage P.1] (NR-54982), hCoV-19/USA/PHC658/2021 [lineage B.1.617.2] (NR-55611). Unless otherwise stated, infection assays were performed with USA-WA1/2020. To propagate each virus strain, sub-confluent monolayers of Vero E6 cells were inoculated with the clinical isolates (MOI < 0.01) and grown for 72 h, at which time significant cytopathic effect was observed for all strains. Culture supernatants were removed, centrifuged 10 min at 1,000 × g, and stored in aliquots at −80°C. To determine titer, focus forming assays were performed on the culture supernatant (assay described in detail below). Substantial differences were noted in the focus phenotypes of these five strains.

#### Lipidomics — Infection

293T-ACE2 cells were seeded at 70% cell density and allowed to grow overnight. Cells were then inoculated with USA-WA1/2020 (MOI = 5) for 1 hr at 37°C in 2% FBS DMEM, rocking gently every 15 minutes. After 1 hr, infection media was removed and replaced with normal 10% DMEM. Cellular lipids were extracted 24 hr after infection.

#### Lipidomics — Transfection

Plasmids containing Strep-tagged viral proteins were generously provided by the Krogan lab at UCSF, and have been described previously (*59*). Wild-type 293T cells were seeded in 6cm dishes and transfected with varying amounts viral plasmids (based on optimal expression for each plasmid, see Table S1), as well as a PLVX empty vector control, using Lipofectamine 3000 (ThermoFisher Scientific) as per manufacturer’s instructions. Transfection media was carefully removed 6 hours after addition and replaced with DMEM. Each condition was repeated in biological quintuplicate. Cellular lipids were extracted 48 hr after transfection.

#### Lipidomics — Lipid Extraction

Cells were washed with PBS and resuspended in a 2 : 1 : 0.75 mixture of chloroform : methanol : water, and 10 μL of an internal standard cocktail (Avanti EquiSPLASH) was added. Extracts were left for one hour at 4 °C, then the layers were separated by centrifugation (3,000xg for 10 minutes), and the chloroform layer was moved to a fresh tube. 2 mL fresh chloroform was added to the aqueous layer, mixed, left for one hour at 4 °C, separated by centrifugation, and then added to the first chloroform layer. The combined chloroform layers were dried under a stream of nitrogen. These dried extracts were frozen at −80 °C and sent to PNNL on dry ice.

#### Lipidomics — LC-MS/MS analysis and lipid identification

LC-MS/MS parameters were established and identifications were conducted as previously described (*60*). A Waters Aquity UPLS H class system interfaced with a Velos-ETD Orbitrap mass spectrometer was used for LC-ESI-MS/MS analyses. Briefly, lipid extracts were dried under vacuum, dissolved in a solution of 10 μL chloroform plus 540 μL of methanol, and 10 μL were injected onto a reverse-phase Waters CSH column (3.0 mmx 150 mm × 1.7 μm particle size), and lipids were separated over a 34-minute gradient (mobile phase A: ACN/H2O (40:60) containing 10 mM ammonium acetate; mobile phase B: ACN/IPA (10:90) containing 10 mM ammonium acetate) at a flow rate of 250 μL/min. Samples were analyzed in both positive and negative mode, using higher-energy collision dissociation and collision-induced dissociation to induce fragmentation. Lipid identifications were made using previously outlined fragment ions (*60*). The LC-MS/MS raw data files were analyzed using LIQUID (*60*), and then all identifications were manually validated by examining the fragmentation spectra for diagnostic and fragment ions corresponding to lipid acyl chains. Identifications were further validated by examining the precursor ion isotopic profile and mass measurement error, extracted ion chromatogram, and retention time for each identified lipid species. To facilitate quantification of lipids, a reference database for lipids identified from the MS/MS data was created, and features from each analysis were then aligned to the reference database based on their m/z, and retention time using MZmine 2 (*61*). Aligned features were manually verified, and peak apex-intensity values were reported for statistical analysis.

#### Lipidomics — QC, normalization, and statistical comparison methods

Lipidomics data were collected in positive and negative ionization mode and analyzed using R. Each ionization mode datasets was normalized using an IS specific to the respective ionization mode. We required that an IS be quantified for every sample to be considered for normalization purposes. Further, normalization factors should not be related to the biological groups being compared to avoid the potential introduction of bias into the data. Thus, for each ionization mode, we evaluated all IS normalization candidates and 1) conducted a test for a difference in mean normalization factors (IS values) by group (Mock vs Virus) and 2) calculated the coefficient of variation (CV) of IS values. The IS showing no evidence of a difference in values by group (p-value > 0.5) and with the minimum CV was selected for normalization. The IS ‘15:0-18:1(d7) PC_IS’ was selected based on the above criteria for both positive and negative ionization data and was used as the normalization factor (log2(abundance/IS abundance)) in both datasets, with a mean CV if 25.8% over the two ionization mode datasets. A one-way analysis of variance (ANOVA) was run on each lipid. The resulting p-values were adjusted for multiple comparisons within each lipid using the Benjamini-Hochberg multiple test correction (*62*).

#### Lipid droplet immunofluorescence — Infection

293T-ACE2 or Caco2 cells were seeded at 70% cell density in 24-well plates and allowed to grow overnight. Cells were then inoculated with USA-WA1/2020 (MOI = 1) for 1 hr at 37°C in 2% FBS DMEM, rocking gently every 15 minutes. After 1 hr, infection media was removed and replaced with normal 10% DMEM (or 20% DMEM, for Caco2 cells). Cells were fixed 8 hours, 24 hours, or 48 hours after infection in 4% PFA.

#### Lipid droplet immunofluorescence — Imaging

After fixation, cells were washed three times with PBS, permeabilized with 0.01% digitonin in PBS for thirty minutes, and blocked with 5% Normal Goat Serum in PBS. Cells were stained overnight with an anti-dsRNA antibody diluted 1:50 in blocking buffer. Cells were washed three times with PBS and then stained with an A647 secondary antibody for 1 hr. Cells were then stained with 1 μg/mL BODIPY 493/503 in PBS for 15 minutes, and then 1x DAPI for 10 minutes. Cells were imaged on a Zeiss LSM 980 Laser-Scanning 3-channel confocal microscope with Airyscan.2.

#### Lipid droplet immunofluorescence — Image analysis

Pearson’s correlation coefficients were measured from 2D projections of z-stacks in Cellprofiler (*63*). Lipid were counted and their sizes, in number of pixels, were measured, using a Cellprofiler pipeline.

#### Cytotoxicity screening

293T-ACE2 and Caco2 cells were seeded in 96-well plates. The next day they were treated with six 5-fold dilutions of each compound, starting from 50 μM. Each condition was tested in triplicate. After 72 hours of compound treatment, cytotoxicity was assessed using resazurin, which is converted into fluorescent resarufin by cells with active oxidative metabolism (*65*). Resazurin was added to a concentration of 0.15 mg/mL and cells were left at 37 °C for 4 hours, and then fluorescence intensity was measured using a BMG CLARIOstar fluorescence plate reader with 560 nm excitation/590 nm emission.

#### Single concentration screen for replication inhibition (all strains of SARS-CoV-2)

The highest concentration for each inhibitor that did not cause cytotoxicity was selected for this assay. For most described inhibitors 10 μM was used, except for MAF (100 μM), and remdesivir (2 μM). Each cell line (Caco2 or 293T-ACE2) was seeded in 96-well plates at a density of 10,000 cells per well and treated overnight with each inhibitor prior to infection with SARS-CoV-2 with an MOI of 0.1. The infection was continued for 48 hours. To quantify viral production, focus-forming assays were performed on the supernatants, described in detail below.

#### Pseudovirus lentivirus production

293T cells were seeded at 2 million cells/dish in 6cm TC-treated dishes. The following day, cells were transfected as described above with lentivirus packaging plasmids, SARS-CoV-2 S plasmid, and IzGreen reporter plasmid (*66*). After transfection, cells were incubated at 37 °C for 60 hours. Viral media was harvested, filtered with a 0.45 μm filter, then frozen before use. Virus transduction capability was then determined by fluorescence using a BZ-X700 all-in-one fluorescent microscope (Keyence), and a 1:16 dilution of viral stocks was found to be optimal for neutralization assays.

#### Pseudovirus entry assay

Neutralization protocol was based on previously reported experiments with the SARS-CoV-2 S pseudotyped lentivirus (*66*). 293T-ACE2 cells were seeded on tissue-culture-treated, poly-lysine treated 96-well plates at a density of 10,000 cells per well. Cells were allowed to grow overnight at 37 °C, and then treated with selected inhibitors as described above for live virus infection. Lz-Green SARS-CoV-2 S pseudotyped lentivirus was added to 293T-ACE2 cells treated with 5 μg/mL polybrene and incubated for 48 hours before imaging. Cells were fixed with 4 % PFA for 1 hour at room temperature, incubated with DAPI for 10 minutes at room temperature, and imaged with BZ-X700 all-in-one fluorescent microscope (Keyence). Estimated area of DAPI and GFP fluorescent pixels was calculated with built-in BZ-X software (Keyence). There were five biological replicates for each condition, and the biggest outlier was removed from analysis due to inherent variability in the assay.

#### Measurement of compound EC_50_

Compounds from the single concentration screen that showed efficacy against SARS-CoV-2 replication were tested to measure compound EC50. The cell line of interest (293T-ACE2 or Caco2) was seeded in 96-well plates at a density of 10,000 cells per well, and treated overnight with 2-fold dilutions of each compound, starting from 50 μM for Atglistatin, PF04620110, GSK2194069, and CAY10499, and starting at 1 μM for Orlistat. Each condition was tested in quadruplicate. The next day cells were infected as described above, and the infection was continued for 48 hours, and then the supernatants were used in a focus forming assay, as described below.

#### Focus forming assay

Vero E6 cells were seeded in a 96-well plate at a density of 20,000 cells per well. The next day, supernatants from infected Caco2 or 293T-ACE2 cells were diluted by adding 225 μL dilution media (Opti-MEM, 2% FBS, 1% pen-strep, 1% non-essential amino acids) to a U-bottom 96-well plate, and then 25 μL of virus-infected supernatant. Further dilutions were made in the same manner, if so desired. Media from the Vero E6 cells was removed and 25 μL diluted virus was added to each well. Vero E6 cells were inoculated for 1 hour at 37 °C/5% CO2 with occasional rocking. After 1 hour, 125 μL of overlay media (0.01 mg/mL methylcellulose in dilution media) was added to each well. Plates were incubated at 37 °C for 24 hours. Overlay media was removed, and replaced with 4% PFA. Plate and lid were saturated in 4% PFA for at least 1 hour at room temperature and removed from the BSL-3. PFA was washed off by gently immersing the plate in a vat of deionized water. Plates were permeabilized in perm buffer (0.1% saponin, 0.1% BSA in PBS) for 30 minutes, then incubated with 50 μL primary antibody (alpaca anti-SARS-CoV-2 serum, diluted 1:5,000 in perm buffer) for either 2hr room temperature or overnight at 4 °C. Antibody was removed and plates were washed 3 × 5 minutes with 200 μL/well PBST (0.1% tween in PBS). Plates were incubated with 50 μL secondary antibody (anti-llama HRP, goat IgG) for either 2hr room temperature or overnight at 4 °C. Antibody was removed and plates were washed 3 × 5 minutes with 200 μL/well PBST. Plates were stained with 50 μL/well TrueBlue peroxidase substrate for 30 minutes. Foci were imaged on an ImmunoSpot S6 Macro ELISPOT imager, and then counted using the Viridot R package (*67*).

#### Quantification and Statistical Analysis

EC_50_ values were calculated using the Hill equation in the R software package. Unless otherwise stated, P values are from one-way ANOVA tests without adjustments for multiple comparisons, with P < 0.05 considered statistically significant.

## Data and Code Availability

The raw lipidomics datasets generated during this study have been deposited and will be available at ftp://massive.ucsd.edu/MSV000087944/. Summaries of fold change changes and p-values are provided in Supplementary Data 1 (live virus lipidomics) and Supplementary Data 2 (viral protein lipidomics).

## Acknowledgments

This work was supported by the National Institutes of Health (1RO1AI141549) and the Pulmonary & Critical Care training grant, “Multidisciplinary Research Training in Pulmonary Medicine” (T32HL083808). Lipidomics analyses were performed in the Environmental Molecular Sciences Laboratory, a national scientific user facility sponsored by the Department of Energy (DOE) Office of Biological and Environmental Research located at the Pacific Northwest National Laboratory (PNNL). PNNL is a multiprogram national laboratory operated by Battelle for the DOE under Contract DE-AC05-76RLO 1830. J.E.K and L.M.B were supported by Laboratory Directed Research and Development Program at PNNL, and the National Institute of Environmental Health Sciences grant U2CES030170. PNNL is a multiprogram national laboratory operated by Battelle for the U.S. Department of Energy under contract DE-AC05-76RLO 1830.

## Author contributions

Author contributions Conceptualization: FGT, SEF, JEK, and HCL; Methodology, formal analysis, and investigation: SEF, JEK, HCL, TAB, JW, LMB; Writing — original draft: SEF; Writing — review and editing: all authors; Visualization: SEF and J-YL; Supervision: FGT; Project administration: FGT; Fund acquisition: FGT, CS, JEK, and TOM.

## Materials & Correspondence

Correspondence and material requests should be addressed to Dr. Fikadu Tafesse.

## References

1. Wolff G, Limpens RWAL, Zevenhoven-Dobbe JC, Laugks U, Zheng S, Jong AWMd, et al. A molecular pore spans the double membrane of the coronavirus replication organelle. Science. 2020;369:1395–8.

2. Klein S, Cortese M, Winter SL, Wachsmuth-Melm M, Neufeldt CJ, Cerikan B, et al. SARS-CoV-2 structure and replication characterized by in situ cryo-electron tomography. Nat Commun. 2020;11(1):5885.

3. Welsch S, Miller S, Romero-Brey I, Merz A, Bleck CK, Walther P, et al. Composition and three-dimensional architecture of the dengue virus replication and assembly sites. Cell Host Microbe. 2009;5(4):365–75.

4. Shulla A, Randall G. (+) RNA virus replication compartments: a safe home for (most) viral replication. Curr Opin Microbiol. 2016;32:82–8.

5. Cortese M, Goellner S, Acosta EG, Neufeldt CJ, Oleksiuk O, Lampe M, et al. Ultrastructural Characterization of Zika Virus Replication Factories. Cell Rep. 2017;18(9):2113–23.

6. Leier HC, Weinstein JB, Kyle JE, Lee JY, Bramer LM, Stratton KG, et al. A global lipid map defines a network essential for Zika virus replication. Nat Commun. 2020;11(1):3652.

7. Melo C, Delafiori J, Dabaja MZ, de Oliveira DN, Guerreiro TM, Colombo TE, et al. The role of lipids in the inception, maintenance and complications of dengue virus infection. Sci Rep. 2018;8(1):11826.

8. Kimhofer T, Lodge S, Whiley L, Gray N, Loo RL, Lawler NG, et al. Integrative Modeling of Quantitative Plasma Lipoprotein, Metabolic, and Amino Acid Data Reveals a Multiorgan Pathological Signature of SARS-CoV-2 Infection. J Proteome Res. 2020;19(11):4442–54.

9. Shen B, Yi X, Sun Y, Bi X, Du J, Zhang C, et al. Proteomic and Metabolomic Characterization of COVID-19 Patient Sera. Cell. 2020.

10. Masana L, Correig E, Ibarretxe D, Anoro E, Arroyo JA, Jerico C, et al. Low HDL and high triglycerides predict COVID-19 severity. Sci Rep. 2021;11(1):7217.

11. Nguyen M, Bourredjem A, Piroth L, Bouhemad B, Jalil A, Pallot G, et al. High plasma concentration of non-esterified polyunsaturated fatty acids is a specific feature of severe COVID-19 pneumonia. Sci Rep. 2021;11(1):10824.

12. Richardson S, Hirsch JS, Narasimhan M, Crawford J, McGinn T, Davidson K. Presenting Characteristics, Comorbidities, and Outcomes Among 5700 Patients Hospitalized With COVID-19 in the New York City Area. Journal of the American Medican Association. 2020;323(20):2052–9.

13. Bligh EG, Dyer WJ. A rapid method of total lipid extraction and purification. Can J Biochem Physiol. 1959;37(8):911–7.

14. Snijder EJ, Limpens R, de Wilde AH, de Jong AWM, Zevenhoven-Dobbe JC, Maier HJ, et al. A unifying structural and functional model of the coronavirus replication organelle: Tracking down RNA synthesis. PLoS Biol. 2020;18(6):e3000715.

15. Goldsmith CS, Tatti KM, Ksiazek TG, Rollin PE, Comer JA, Lee WW, et al. Ultrastructural Characterization of SARS Coronavirus. Emerging Infectious Diseases. 2004;10(2):320–7.

16. Mohan J, Wollert T. Membrane remodeling by SARS-CoV-2 - double-enveloped viral replication. Fac Rev. 2021;10:17.

17. Angelini MM, Akhlaghpour M, Neuman BW, Buchmeier MJ. Severe acute respiratory syndrome coronavirus nonstructural proteins 3, 4, and 6 induce double-membrane vesicles. mBio. 2013;4(4).

18. Hagemeijer MC, Monastyrska I, Griffith J, van der Sluijs P, Voortman J, van Bergen en Henegouwen PM, et al. Membrane rearrangements mediated by coronavirus nonstructural proteins 3 and 4. Virology. 2014;458-459:125–35.

19. Zhou H, Ferraro D, Zhao J, Hussain S, Shao J, Trujillo J, et al. The N-terminal region of severe acute respiratory syndrome coronavirus protein 6 induces membrane rearrangement and enhances virus replication. J Virol. 2010;84(7):3542–51.

20. Banerjee AK, Blanco MR, Bruce EA, Honson DD, Chen LM, Chow A, et al. SARS-CoV-2 Disrupts Splicing, Translation, and Protein Trafficking to Suppress Host Defenses. Cell. 2020.

21. Angeletti S, Benvenuto D, Bianchi M, Giovanetti M, Pascarella S, Ciccozzi M. COVID-2019: The role of the nsp2 and nsp3 in its pathogenesis. J Med Virol. 2020.

22. Nelson CA, Pekosz A, Fremont DH. Structure and Intracellular Targeting of the SARS-Coronavirus Orf7a Acessory Protein. Structure. 2005;13:75–85.

23. Schaecher SR, Diamond MS, Pekosz A. The transmembrane domain of the severe acute respiratory syndrome coronavirus ORF7b protein is necessary and sufficient for its retention in the Golgi complex. J Virol. 2008;82(19):9477–91.

24. Chen CC, Kruger J, Sramala I, Hsu HJ, Henklein P, Chen YM, et al. ORF8a of SARS-CoV forms an ion channel: experiments and molecular dynamics simulations. Biochim Biophys Acta. 2011;1808(2):572–9.

25. Bianchi M, Borsetti A, Ciccozzi M, Pascarella S. SARS-Cov-2 ORF3a: Mutability and function. Int J Biol Macromol. 2021;170:820–6.

26. Meier C, Aricescu AR, Assenberg R, Aplin RT, Gilbert RJ, Grimes JM, et al. The crystal structure of ORF-9b, a lipid binding protein from the SARS coronavirus. Structure. 2006;14(7):1157–65.

27. Chen X, Wang K, Xing Y, Tu J, Yang X, Zhao Q, et al. Coronavirus membrane-associated papain-like proteases induce autophagy through interacting with Beclin1 to negatively regulate antiviral innate immunity. Protein Cell. 2014;5(12):912–27.

28. Cottam EM, Maier HJ, Manifava M, Vaux LC, Chandra-Schoenfelder P, Gerner W, et al. Coronavirus nsp6 proteins generate autophagosomes from the endoplasmic reticulum via an omegasome intermediate. Autophagy. 2011;7(11):1335–47.

29. Yue Y, Nabar NR, Shi CS, Kamenyeva O, Xiao X, Hwang IY, et al. SARS-Coronavirus Open Reading Frame-3a drives multimodal necrotic cell death. Cell Death Dis. 2018;9(9):904.

30. Ren Y, Shu T, Wu D, Mu J, Wang C, Huang M, et al. The ORF3a protein of SARS-CoV-2 induces apoptosis in cells. Cell Mol Immunol. 2020;17(8):881–3.

31. Ye Z, Wong CK, Li P, Xie Y. A SARS-CoV protein, ORF-6, induces caspase-3 mediated, ER stress and JNK-dependent apoptosis. Biochim Biophys Acta. 2008;1780(12):1383–7.

32. Tan YX, Tan TH, Lee MJ, Tham PY, Gunalan V, Druce J, et al. Induction of apoptosis by the severe acute respiratory syndrome coronavirus 7a protein is dependent on its interaction with the Bcl-XL protein. J Virol. 2007;81(12):6346–55.

33. Singh R, Kaushik S, Wang Y, Xiang Y, Novak I, Komatsu M, et al. Autophagy regulates lipid metabolism. Nature. 2009;458(7242):1131–5.

34. Crimi M, Esposti MD. Apoptosis-induced changes in mitochondrial lipids. Biochim Biophys Acta. 2011;1813(4):551–7.

35. Welte MA, Gould AP. Lipid droplet functions beyond energy storage. Biochim Biophys Acta Mol Cell Biol Lipids. 2017;1862(10 Pt B):1260–72.

36. Miyanari Y, Atsuzawa K, Usuda N, Watashi K, Hishiki T, Zayas M, et al. The lipid droplet is an important organelle for hepatitis C virus production. Nat Cell Biol. 2007;9(9):1089–97.

37. Roingeard P, Hourioux C. Hepatitis C virus core protein, lipid droplets and steatosis. J Viral Hepat. 2008;15(3):157–64.

38. Heaton NS, Randall G. Dengue Virus-Induced Autophagy Regulates Lipid Metabolism. Cell Host & Microbe. 2010;8:422–32.

39. Qiu B, Simon MC. BODIPY 493/503 Staining of Neutral Lipid Droplets for Microscopy and Quantification by Flow Cytometry. Bio Protoc. 2016;6(17).

40. Kumar P, Nagarajan A, Uchil PD. Analysis of Cell Viability by the alamarBlue Assay. Cold Spring Harb Protoc. 2018;2018(6).

41. Case JB, Bailey AL, Kim AS, Chen RE, Diamond MS. Growth, detection, quantification, and inactivation of SARS-CoV-2. Virology. 2020;548:39–48.

42. Hardwicke MA, Rendina AR, Williams SP, Moore ML, Wang L, Krueger JA, et al. A human fatty acid synthase inhibitor binds beta-ketoacyl reductase in the keto-substrate site. Nat Chem Biol. 2014;10(9):774–9.

43. Hadváry P, Lengsfeld H, Wolfer H. Inhibition of pancreatic lipase*in vitro*by the covalent inhibitor tetrahydrolipstatin. Biochem J. 1988;256:357–61.

44. Kridel SJ, Axelrod F, Rozenkrantz N, Smith JW. Orlistat Is a Novel Inhibitor of Fatty Acid Synthase with Antitumor Activity. Cancer Research. 2004;64:2070–5.

45. Dow RL, Li JC, Pence MP, Gibbs EM, LaPerle JL, Litchfield J, et al. Discovery of PF-04620110, a Potent, Selective, and Orally Bioavailable Inhibitor of DGAT-1. ACS Med Chem Lett. 2011;2(5):407–12.

46. Kraemer FB, Shen W-J. Hormone-sensitive lipase: control of intracellular tri-(di-)acylglycerol and cholesterol ester hydrolysis. Journal of Lipid Research. 2002;43(10):1585–94.

47. Muccioli GG, Labar G, Lambert DM. CAY10499, a novel monoglyceride lipase inhibitor evidenced by an expeditious MGL assay. Chembiochem. 2008;9(16):2704–10.

48. Mayer N, Schweiger M, Romauch M, Grabner GF, Eichmann TO, Fuchs E, et al. Development of small-molecule inhibitors targeting adipose triglyceride lipase. Nat Chem Biol. 2013;9(12):785–7.

49. Bleasdale JE, Thakur NR, Gremban RS, Bundy GL, Fitzpatrick FA, Smith RJ, et al. Selective Inhibition of Receptor-Coupled Phospholipse-C-Dependent Processes in Human Platelets and Polymorphonuclear Neutrophils. The Journal of Pharmacology and Experimental Therapeutics. 1990;255(2):756–68.

50. Graham MS, Sudre CH, May A, Antonelli M, Murray B, Varsavsky T, et al. Changes in symptomatology, reinfection, and transmissibility associated with the SARS-CoV-2 variant B.1.1.7: an ecological study. Lancet Public Health. 2021;6:e335–45.

51. Cele S, Gazy I, Jackson L, Hwa SH, Tegally H, Lustig G, et al. Escape of SARS-CoV-2 501Y.V2 from neutralization by convalescent plasma. Nature. 2021;593(7857):142–6.

52. Wang P, Casner RG, Nair MS, Wang M, Yu J, Cerutti G, et al. Increased resistance of SARS-CoV-2 variant P.1 to antibody neutralization. Cell Host Microbe. 2021;29(5):747–51 e4.

53. Kupferschmidt K. Fast-spreading U.K. virus variant raises alarms. Science. 2021;371(6524):9–10.

54. Tegally H, Wilkinson E, Giovanetti M, Iranzadeh A, Fonseca V, Giandhari J, et al. Emergence and rapid spread of a new severe acute respiratory syndrome-related coronavirus 2(SARS-CoV-2) ilneage with multiple spike mutations in South Africa. medRxiv. 2020.

55. Faria NR, Mellan TA, Whittaker C, Claro IM, Candido DdS, Mishra S, et al. Genomics and epidemiology of the P.1 SARS-CoV-2 lineage in Manaus, Brazil. Science. 2021;372:815–21.

56. Liu C, Ginn HM, Dejnirattisai W, Supasa P, Wang B, Tuekprakhon A, et al. Reduced neutralization of SARS-CoV-2 B.1.617 by vaccine and convalescent serum. Cell. 2021.

57. Hadfield J, Megill C, Bell SM, Huddleston J, Potter B, Callender C, et al. Nextstrain: real-time tracking of pathogen evolution. Bioinformatics. 2018;34(23):4121–3.

58. Sagulenko P, Puller V, Neher RA. TreeTime: Maximum-likelihood phylodynamic analysis. Virus Evol. 2018;4(1):vex042.

59. Gordon, D. E.; Jang, G. M.; Bouhaddou, M.; Xu, J.; Obernier, K.; White, K. M.; O’Meara, M. J.; Rezelj, V. V.; Guo, J. Z.; Swaney, D. L.; Tummino, T. A.; Huettenhain, R.; Kaake, R. M.; Richards, A. L.; Tutuncuoglu, B.; Foussard, H.; Batra, J.; Haas, K.; Modak, M.; Kim, M.; Haas, P.; Polacco, B. J.; Braberg, H.; Fabius, J. M.; Eckhardt, M.; Soucheray, M.; Bennett, M. J.; Cakir, M.; McGregor, M. J.; Li, Q.; Meyer, B.; Roesch, F.; Vallet, T.; Mac Kain, A.; Miorin, L.; Moreno, E.; Naing, Z. Z. C.; Zhou, Y.; Peng, S.; Shi, Y.; Zhang, Z.; Shen, W.; Kirby, I. T.; Melnyk, J. E.; Chorba, J. S.; Lou, K.; Dai, S. A.; Barrio-Hernandez, I.; Memon, D.; Hernandez-Armenta, C.; Lyu, J.; Mathy, C. J. P.; Perica, T.; Pilla, K. B.; Ganesan, S. J.; Saltzberg, D. J.; Rakesh, R.; Liu, X.; Rosenthal, S. B.; Calviello, L.; Venkataramanan, S.; Liboy-Lugo, J.; Lin, Y.; Huang, X. P.; Liu, Y.; Wankowicz, S. A.; Bohn, M.; Safari, M.; Ugur, F. S.; Koh, C.; Savar, N. S.; Tran, Q. D.; Shengjuler, D.; Fletcher, S. J.; O’Neal, M. C.; Cai, Y.; Chang, J. C. J.; Broadhurst, D. J.; Klippsten, S.; Sharp, P. P.; Wenzell, N. A.; Kuzuoglu, D.; Wang, H. Y.; Trenker, R.; Young, J. M.; Cavero, D. A.; Hiatt, J.; Roth, T. L.; Rathore, U.; Subramanian, A.; Noack, J.; Hubert, M.; Stroud, R. M.; Frankel, A. D.; Rosenberg, O. S.; Verba, K. A.; Agard, D. A.; Ott, M.; Emerman, M.; Jura, N.; von Zastrow, M.; Verdin, E.; Ashworth, A.; Schwartz, O.; d’Enfert, C.; Mukherjee, S.; Jacobson, M.; Malik, H. S.; Fujimori, D. G.; Ideker, T.; Craik, C. S.; Floor, S. N.; Fraser, J. S.; Gross, J. D.; Sali, A.; Roth, B. L.; Ruggero, D.; Taunton, J.; Kortemme, T.; Beltrao, P.; Vignuzzi, M.; Garcia-Sastre, A.; Shokat, K. M.; Shoichet, B. K.; Krogan, N. J., A SARS-CoV-2 protein interaction map reveals targets for drug repurposing. Nature 2020.

60. Kyle, J. E.; Crowell, K. L.; Casey, C. P.; Fujimoto, G. M.; Kim, S.; Dautel, S. E.; Smith, R. D.; Payne, S. H.; Metz, T. O., LIQUID: an-open source software for identifying lipids in LC-MS/MS-based lipidomics data. Bioinformatics 2017, 33 (11), 1744–1746.

61. Pluskal, T.; Castillo, S.; Villar-Briones, A.; Orešič, M., MZmine 2: Modular framework for processing, visualizing, and anlyzing mass spectrometry-based molecular profile data. BMC Bioinformatics 2010, 11 (395), 1–11.

62. Benjamini, Y.; Hochberg, Y., Controlling the False Discovery Rate: A Practical an Powerful Approach to Multiple Testing. Journal of the Royal Statistical Society: Series B (Methodological) 1995, 57, 289–300.

63. McQuin, C.; Goodman, A.; Chernyshev, V.; Kamentsky, L.; Cimini, B. A.; Karhohs, K. W.; Doan, M.; Ding, L.; Rafelski, S. M.; Thirstrup, D.; Wiegraebe, W.; Singh, S.; Becker, T.; Caicedo, J. C.; Carpenter, A. E., CellProfiler 3.0: Next-generation image processing for biology. PLoS Biol 2018, 16 (7), e2005970.

64. Schindelin, J.; Arganda-Carreras, I.; Frise, E.; Kaynig, V.; Longair, M.; Pietzsch, T.; Preibisch, S.; Rueden, C.; Saalfeld, S.; Schmid, B.; Tinevez, J. Y.; White, D. J.; Hartenstein, V.; Eliceiri, K.; Tomancak, P.; Cardona, A., Fiji: an open-source platform for biological-image analysis. Nat Methods 2012, 9 (7), 676–82.

65. Kumar, P.; Nagarajan, A.; Uchil, P. D., Analysis of Cell Viability by the alamarBlue Assay. Cold Spring Harb Protoc 2018, 2018 (6).

66. Crawford, K. H. D.; Eguia, R.; Dingens, A. S.; Loes, A. N.; Malone, K. D.; Wolf, C. R.; Chu, H. Y.; Tortorici, M. A.; Veesler, D.; Murphy, M.; Pettie, D.; King, N. P.; Balazs, A. B.; Bloom, J. D., Protocol and Reagents for Pseudotyping Lentiviral Particles with SARS-CoV-2 Spike Protein for Neutralization Assays. Viruses 2020, 12 (5).

67. Katzelnick, L. C.; Coello Escoto, A.; McElvany, B. D.; Chavez, C.; Salje, H.; Luo, W.; Rodriguez-Barraquer, I.; Jarman, R.; Durbin, A. P.; Diehl, S. A.; Smith, D. J.; Whitehead, S. S.; Cummings, D. A. T., Viridot: An automated virus plaque (immunofocus) counter for the measurement of serological neutralizing responses with application to dengue virus. PLoS Negl Trop Dis 2018, 12 (10), e0006862.

68. Hadfield, J.; Megill, C.; Bell, S. M.; Huddleston, J.; Potter, B.; Callender, C.; Sagulenko, P.; Bedford, T.; Neher, R. A., Nextstrain: real-time tracking of pathogen evolution. Bioinformatics 2018, 34 (23), 4121–4123.

69. Sagulenko, P.; Puller, V.; Neher, R. A., TreeTime: Maximum-likelihood phylodynamic analysis. Virus Evol 2018, 4 (1), vex042.

