## Supplemental Figures for "A global lipid map reveals host dependency factors conserved across SARS-CoV-2 variants"

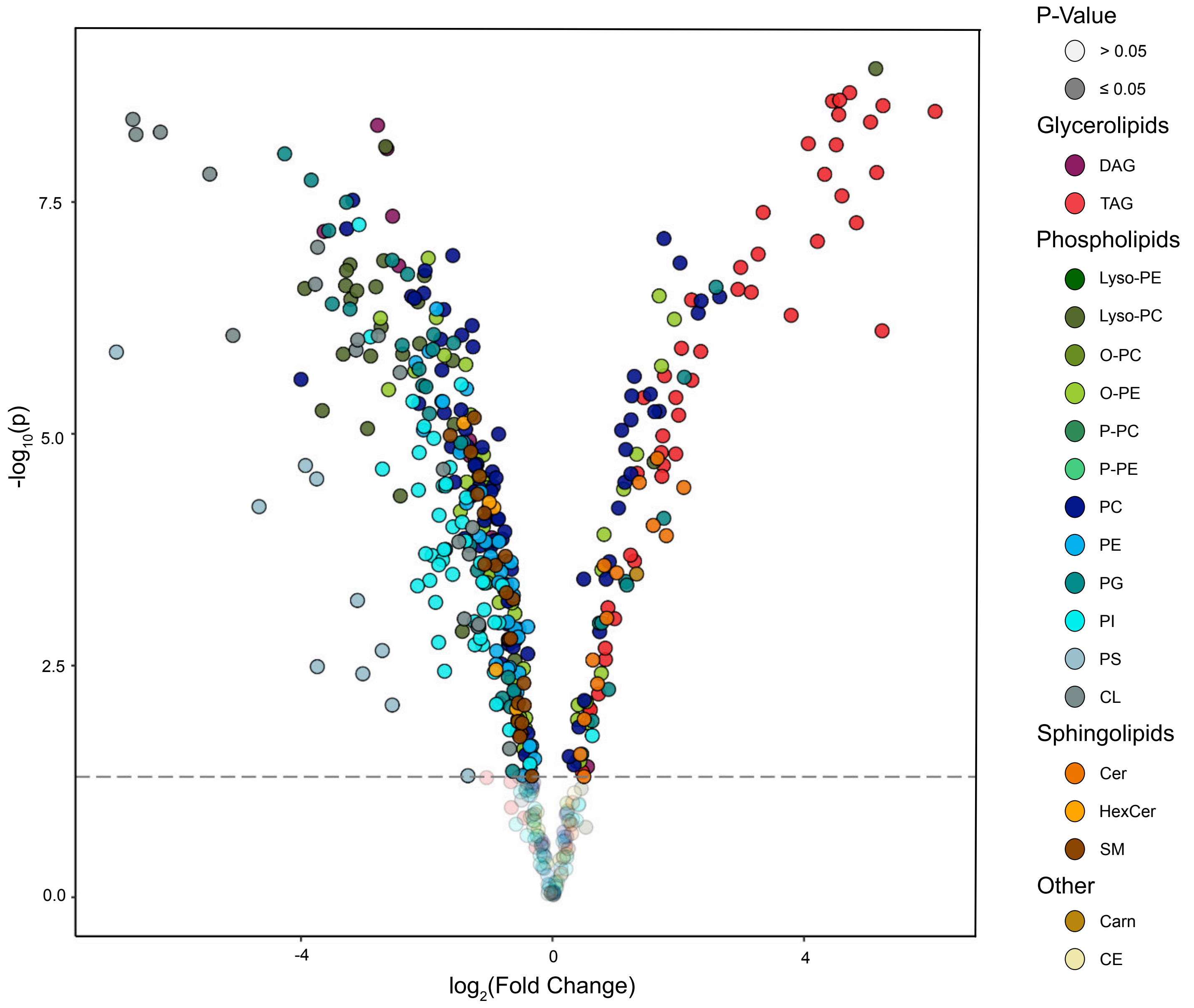


**Fig. S1.**

Volcano plot of individual lipid species detected by LC-ESI-MS/MS. Fold change is calculated relative to abundance in mock-infected controls. Lipid species are colored by class. Abbreviations: DAG = diacylglycerol; TAG = triacylglycerol; Lyso-PC = lysophosphatidylcholine; Lyso-PE = lysophosphatidylethanolamine; O-PC = phosphatidylcholine (ether linked); O-PE = phosphatidylethanolamine (ether linked); P-PC = phosphatidylcholine (plasmologen-linked); P-PE = phosphatidylethanolamine (plasmologen-linked); PC = phosphatidylcholine; PE = phosphatidylethanolamine; PG = phosphatidylglycerol; PI = phosphatidylinositol; PS = phosphatidylserine; CL = cardiolipin; Cer = ceramide; HexCer = hexosylceramide; SM = sphingomyelin; Carn = acylcarnitine; CE = cholesterol ester

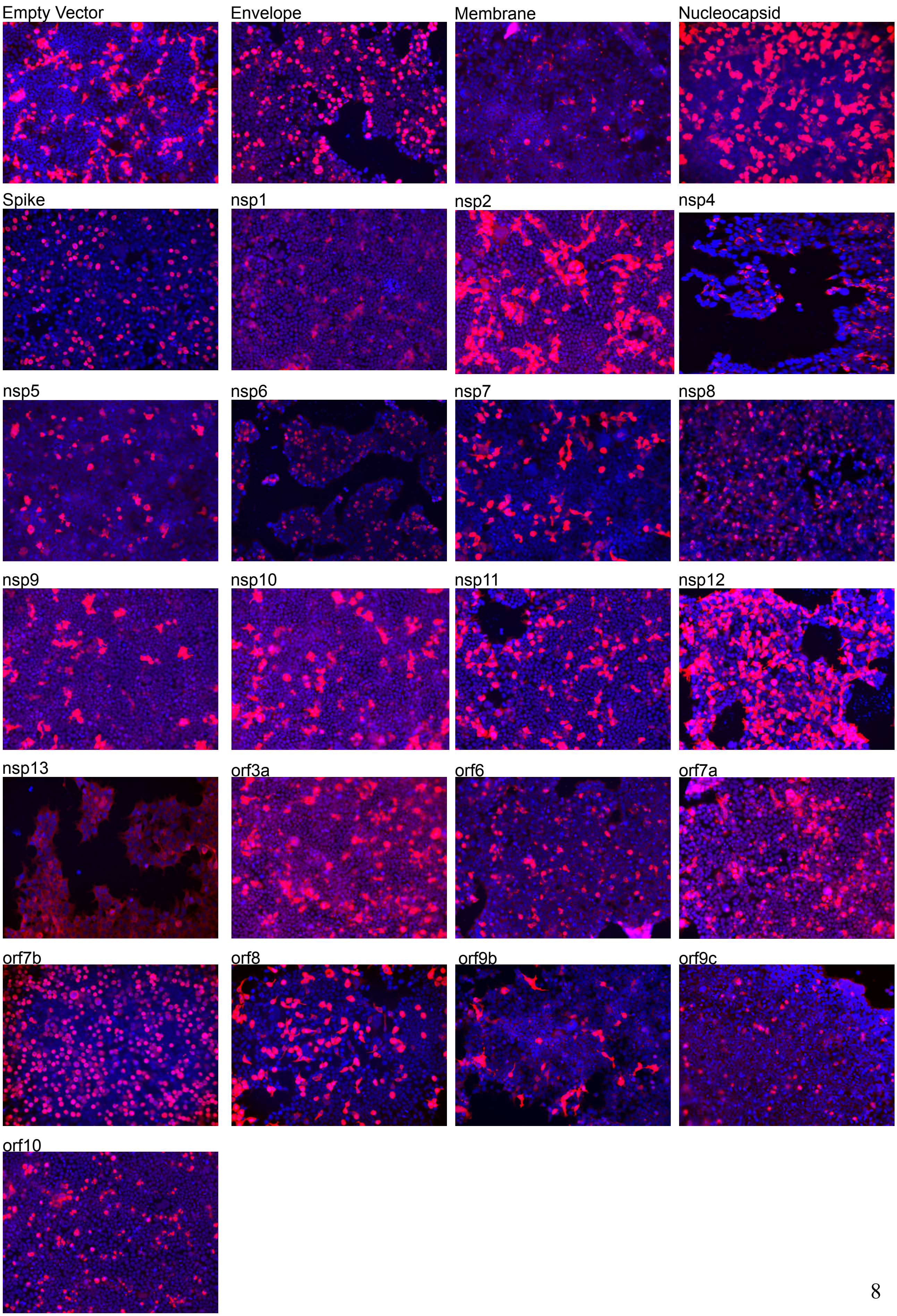
Empty Vector
Envelope
Membrane
Nucleocapsid
Spike
nsp1
nsp2
nsp4
nsp5
nsp6
nsp7
nsp8
nsp9
nsp10
nsp11
nsp12
nsp13
orf3a
orf6
orf7a
orf7b
orf8
orf9b
orf9c
orf10

**Fig. S2.**

Images of HEK293T cells transfected with each viral protein taken on a BZ-X700 all-in-one fluorescent microscope (Keyence) at 20x resolution. Red is anti-Strep-tag immunostaining; blue is DAPI.

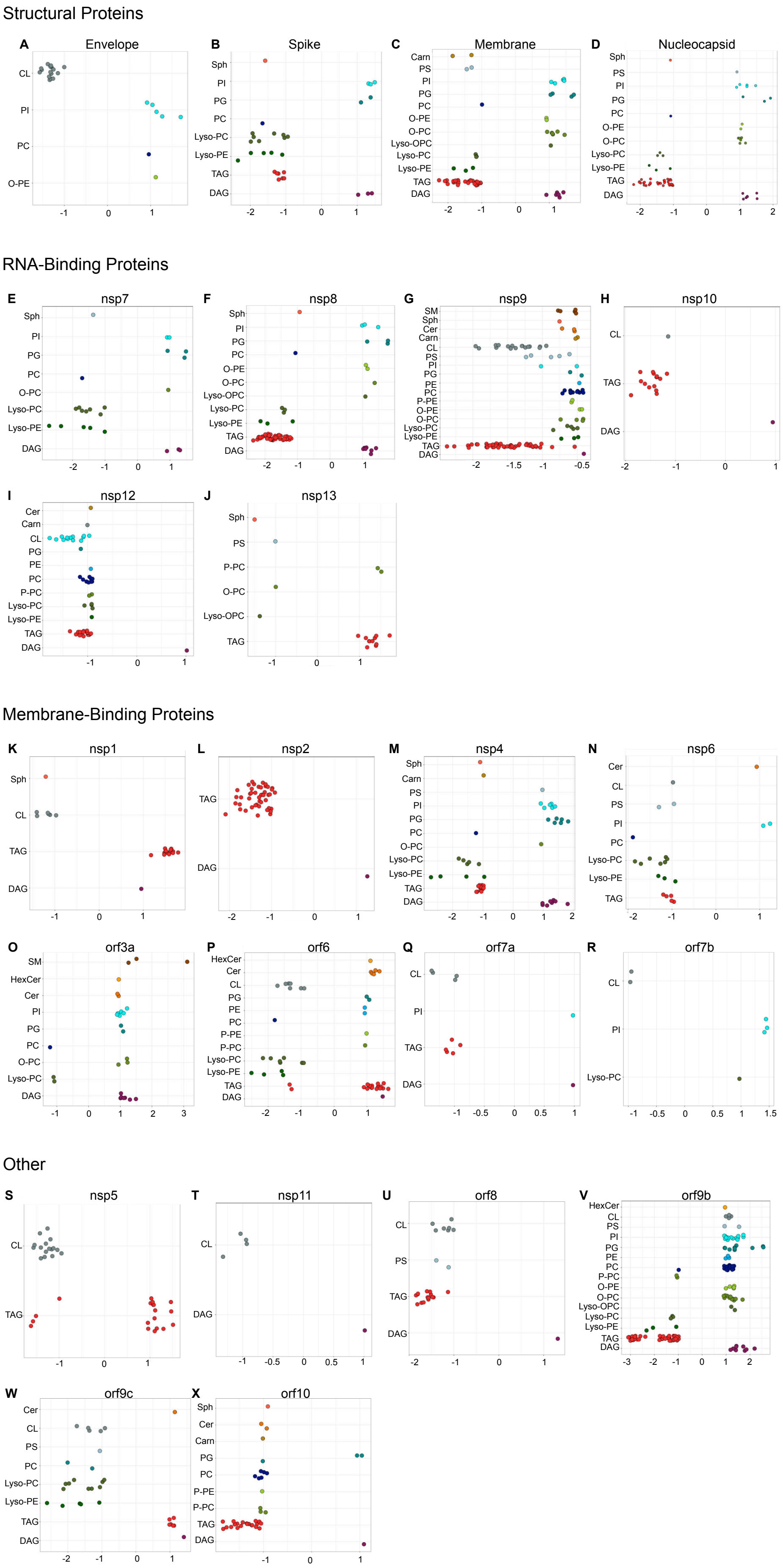
Structural Proteins
A Envelope
CL PI PC O-PE
B Spike
Sph PI PG PC Lyso-PC Lyso-PE TAG DAG
C Membrane
Carn PS PI PG PC O-PE O-PC Lyso-OPC Lyso-PC Lyso-PE TAG DAG
D Nucleocapsid
Sph PS PI PG PC O-PE O-PC Lyso-PC Lyso-PE TAG DAG
RNA-Binding Proteins
E nsp7
Sph PI PG PC O-PC Lyso-PC Lyso-PE DAG
F nsp8
Sph PI PG PC O-PE O-PC Lyso-OPC Lyso-PC Lyso-PE TAG DAG
G nsp9
SM Sph Cer Carn CL PS PI PG PE PC P-PE O-PE O-PC Lyso-PC Lyso-PE TAG DAG
H nsp10
CL TAG DAG
I nsp12
Cer Carn CL PG PE PC P-PC Lyso-PC Lyso-PE TAG DAG
J nsp13
Sph PS P-PC O-PC Lyso-OPC TAG
Membrane-Binding Proteins
K nsp1
Sph CL TAG DAG
L nsp2
TAG DAG
M nsp4
Sph Carn PS PI PG PC O-PC Lyso-PC Lyso-PE TAG DAG
N nsp6
Cer CL PS PI PC Lyso-PC Lyso-PE TAG
O orf3a
SM HexCer Cer PI PG PC O-PC Lyso-PC DAG
P orf6
HexCer Cer CL PG PE PC P-PE P-PC Lyso-PC Lyso-PE TAG DAG
Q orf7a
CL PI TAG DAG
R orf7b
CL PI Lyso-PC
Other
S nsp5
CL TAG
T nsp11
CL DAG
U orf8
CL PS TAG DAG
V orf9b
HexCer CL PS PI PG PE PC P-PC O-PE O-PC Lyso-OPC Lyso-PC Lyso-PE TAG DAG
W orf9c
Cer CL PS PC Lyso-PC Lyso-PE TAG DAG
X orf10
Sph Cer Carn PG PC P-PE P-PC TAG DAG

**Fig. S3.**

For each viral protein, the lipids that changed significantly relative to empty vector were selected. To identify the most influenced pathways, only lipids with a log2(fold change) greater than 0.9 or less than 0.9 were plotted. Log2(fold change) relative to empty vector for each individual lipid species is shown.

(A-D) Structural proteins
(E-J) RNA-binding proteins
(K-R) Membrane-binding proteins
(S-X) Other proteins

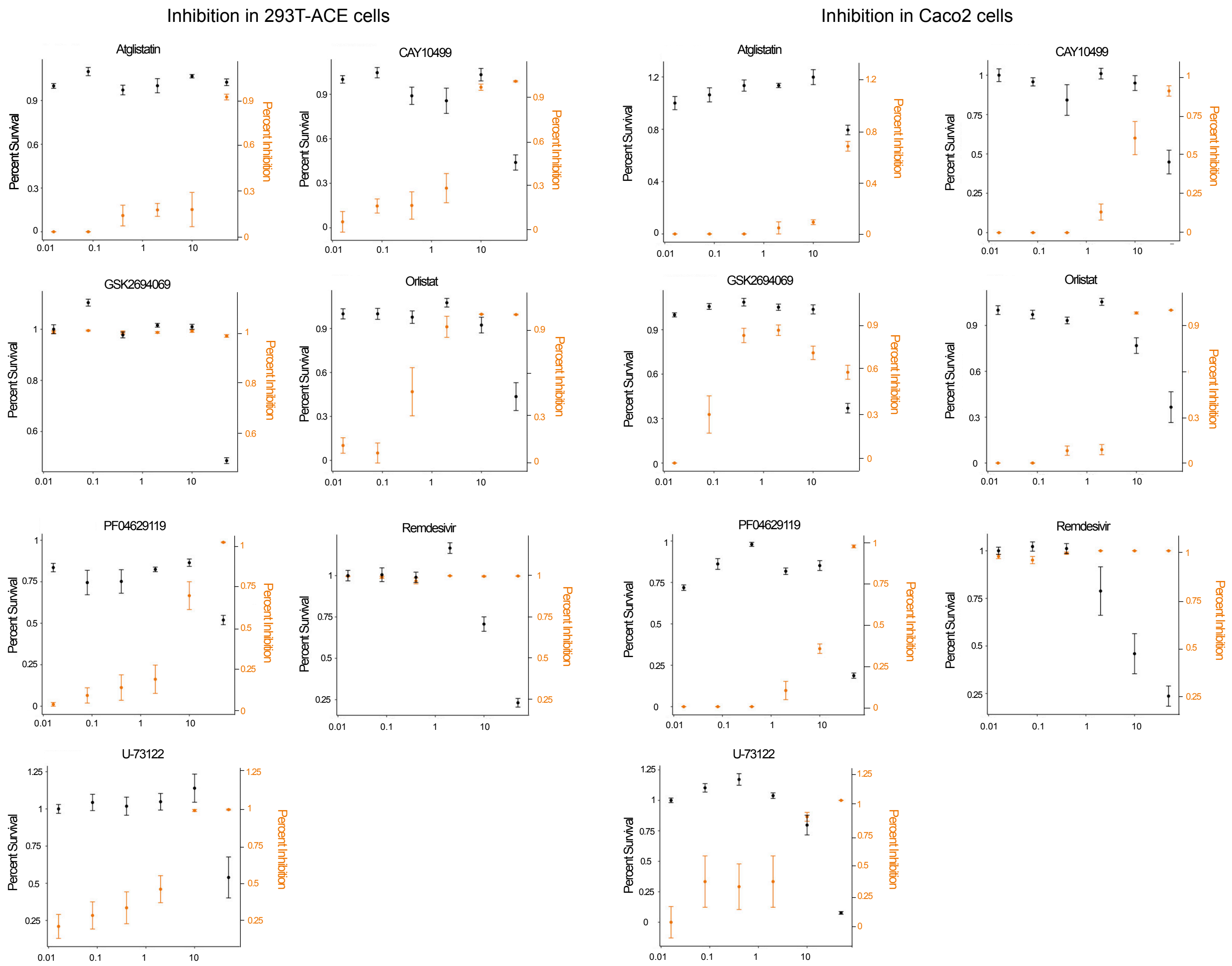


**Fig. S4.**
Percent inhibition/percent survival curves for each compound in 293T-Ace2 cells or Caco2 cells. Percent survival was calculated by dividing the resazurin fluorescence in drug-treated wells by the resazurin fluorescence in vehicle-treated wells. Percent inhibition was calculated by dividing the number of foci in drug-treated wells by the number of foci in vehicle-treated wells. Error bars represent standard error, calculated from three independent experiments.

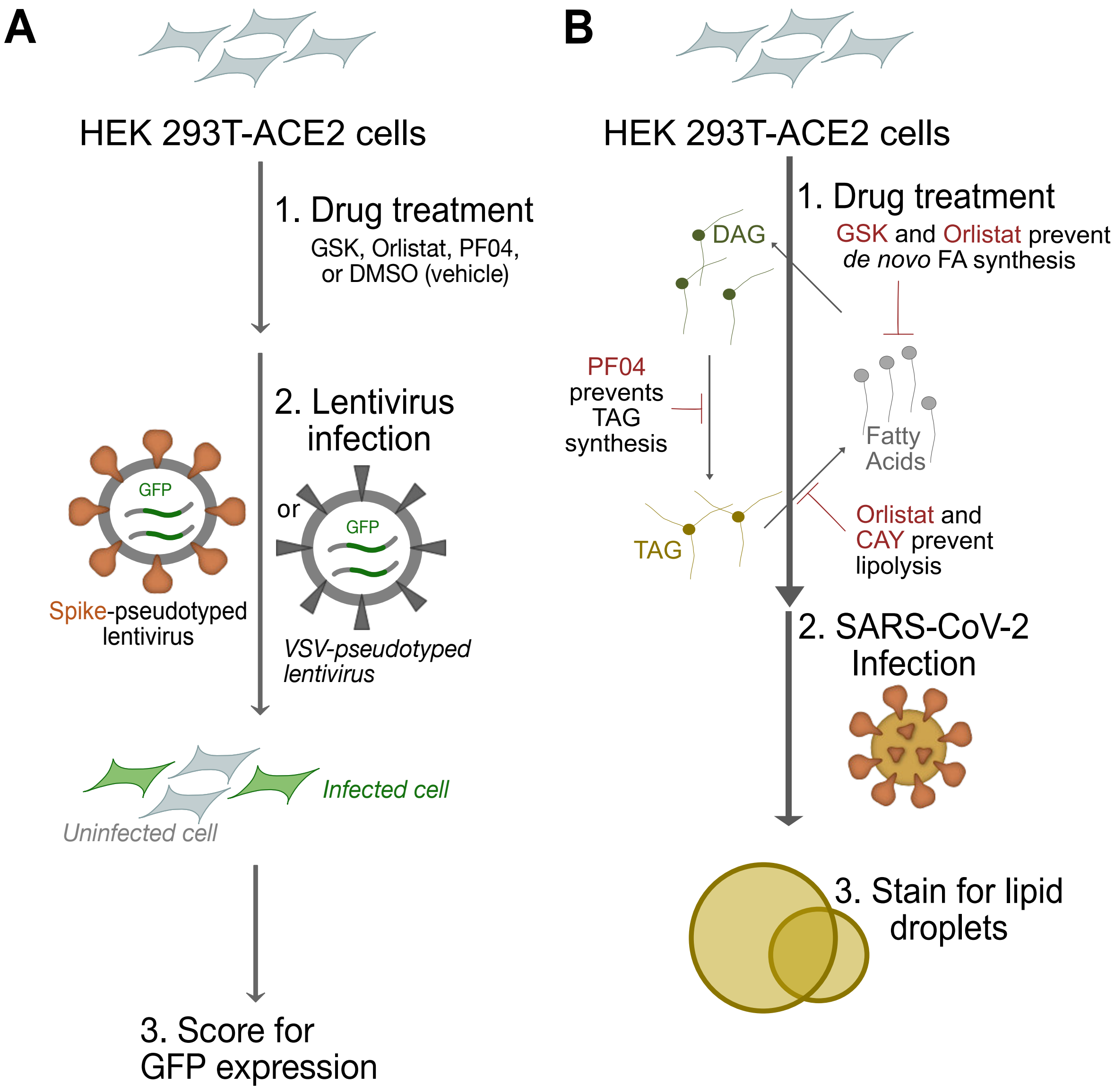


**Fig. S5.**
(**A**) Spike-mediated SARS-CoV-2 entry assay using Spike-pseudotyped lentivirus: experimental scheme. HEK293T-ACE2 cells are treated overnight with 10 µM of selected inhibitors then infected with lentiviruses coated expressing the SARS-CoV-2 Spike protein (or the VSV G protein as a control) and bearing a GFP construct. Successfully infected cells express GFP; GFP fluorescence (normalized to DAPI) is used to quantify Spike-mediated viral uptake (**B**) Experimental design to test the effect of glycerolipid biosynthesis inhibition on SARS-CoV-2 induced lipid droplet formation. HEK293T-ACE2 cells were treated overnight with 10 µM of selected inhibitors and then infected with SARS-CoV-2 (MOI = 1) and fixed after 48 hours. Lipid droplets and infected cells were vidsualized with BODIPY 493/503 and anti-dsRNA immunofluorescence, respectively.

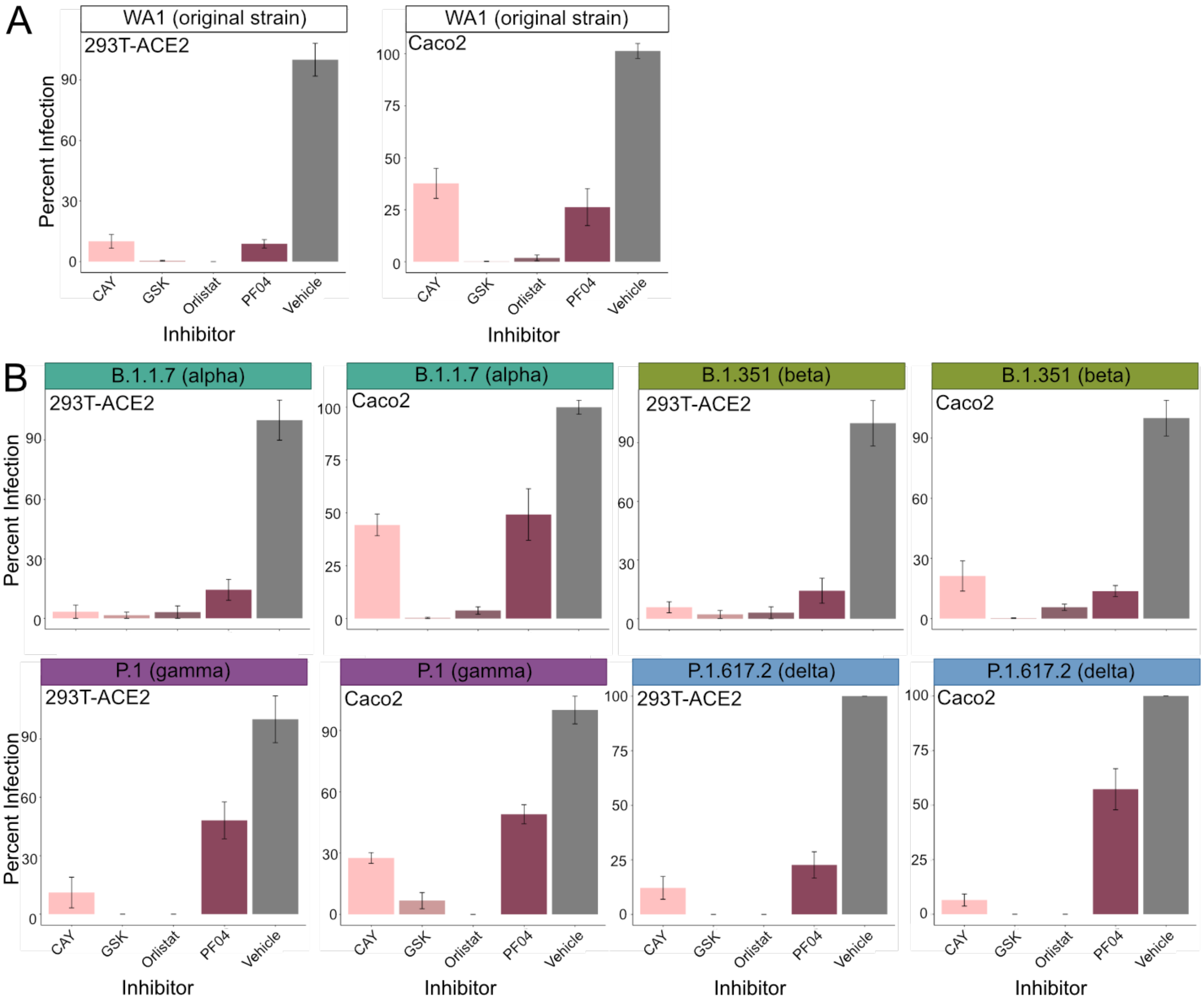


**Fig. S6.**
(**A**) Inhibition of the original SARS-CoV-2 strain by four inhibitors of glycerolipid biosynthesis, each at 10 μM overnight prior to a 48-hour infection (MOI = 0.1). Data are from three independent experiments; data are mean ± SE. (**B**) Inhibition of four variants of concern by four inhibitors of glycerolipid biosynthesis, each at 10 μM overnight prior to a 48-hour infection (MOI = 0.1). Data are from three independent experiments; data are mean ± SE.

| Viral Protein | Plasmid ID | Amount of DNA (μg) |
|---|---|---|
| nsp1 | A01 | 30 |
| nsp2 | A02 | 30 |
| nsp4 | A05 | 30 |
| nsp5 | A06 | 30 |
| nsp7 | A09 | 20 |
| nsp8 | A10 | 30 |
| nsp9 | A11 | 40 |
| nsp10 | A12 | 35 |
| nsp11 | B01 | 30 |
| nsp12 | B02 | 45 |
| nsp13 | B03 | 35 |
| S | B07 | 30 |
| orf3a | B08 | 30 |
| E | B10 | 30 |
| M | B11 | 20 |
| orf6 | B12 | 35 |
| orf7a | C01 | 35 |
| orf7b | C02 | 30 |
| orf8 | C03 | 30 |
| N | C04 | 10 |
| orf9b | C05 | 30 |
| orf9c | C06 | 35 |
| orf10 | C07 | 45 |
| Empty Vector | PLVX | 35 |

**Table S1.**

Amounts of DNA used in viral protein transfection, related to Figure 2 and Figure S2.

**Data S1. (separate file)**

Fold changes and p-values for each lipid observed in the live virus infection experiment

**Data S2. (separate file)**

Fold changes and p-values for each lipid observed in the viral protein transfection experiment
